# Hybrid incompatibility emerges at the one-cell stage in interspecies *Caenorhabditis* embryos

**DOI:** 10.1101/2024.10.19.619171

**Authors:** Jessica Bloom, Rebecca Green, Arshad Desai, Karen Oegema, Scott A. Rifkin

**Author notes:** Lead Contact: Scott A. Rifkin. Corresponding authors: Scott A. Rifkin, Rebecca Green, Karen Oegema.

## Abstract

Intrinsic reproductive isolation occurs when genetic divergence between populations disrupts hybrid development, preventing gene flow and enforcing speciation.^1–4^ Over the past two decades, researchers have identified molecular mechanisms underlying a few dozen cases of hybrid incompatibility in animals.^5^ Much of this work has focused on mismatches in zygotic gene regulation,^6–11^ but other mechanisms have also emerged, including symbiont-driven incompatibilities,^12^ nucleoporin mismatches affecting nuclear-cytoplasmic transport,^13^ and divergence in centromeric or heterochromatic regions and their regulatory proteins which can lead to the inability of the oocyte cytoplasm to segregate sperm-derived chromosomes.^14–19^ Since studies to date have focused on a limited number of species, uncovering mechanisms across diverse taxa will be important to understanding broader patterns of hybrid incompatibility.

Here, we investigate the mechanistic basis of hybrid incompatibility in *Caenorhabditis* nematodes by leveraging the ability of *C. brenneri* females to produce embryos after mating with males from several other species. We find that incompatibilities emerge between fertilization and the onset of zygotic transcription, which begins at the 4-cell stage.^20–23^ In *Caenorhabditis* embryos, as in many animals,^24,25^ sperm deliver chromatin and centrioles into the oocyte.^26–29^ After remaining quiescent during oocyte meiosis, the sperm chromatin acquires a nuclear envelope, and centrioles initiate centrosome formation.^30–32^ Centrosomes remain tethered to the sperm pronucleus, which positions them near the cortex to establish anterior-posterior polarity.^33,34^ We identify two key processes that are destabilized in hybrids: (1) the ability of oocytes to control sperm-derived pronuclear expansion, and (2) successful polar body formation. When sperm pronuclear expansion is delayed, centrosomes detach, which leads to defects in polarity establishment. Hybrid embryos typically experience one or more stochastic failures of early developmental events that accumulate and eventually kill them.

## RESULTS AND DISCUSSION

### Hybrid embryos of *C. brenneri* females and *C. elegans* males exhibit polarity defects prior to zygotic genome activation

In the *Caenorhabditis Elegans* group, most species pairs will not mate. Among those that do, hybrid embryos, with rare exceptions, die during embryogenesis.^35–41^ Notably, within a single cross, the stage at which development arrests can vary widely among individual embryos.^42,43^ Although toxin-antidote systems underlie several incompatibilities between strains within a species,^44–47^ the cellular and molecular bases of interspecies hybrid incompatibility in *Caenorhabditis* remain unknown.

To identify incompatibilities that impair the development of hybrid embryos, we took advantage of the ability of *C. brenneri* females to be fertilized by males from multiple species across the *Elegans* group (**Fig. 1A**; eight fertilizing species highlighted in dark grey). To take advantage of tools available in *C. elegans*, we began by analyzing embryos from *C. brenneri* oocytes fertilized with *C. elegans* sperm. As expected, all hybrid embryos died, compared to 6% (145/2263) and 2% (56/2991) embryonic lethality in intraspecies *C. brenneri* and *C. elegans* embryos, respectively (**Fig. 1B**); the higher intra-specific lethality in *C. brenneri* is expected due to ongoing inbreeding depression in lab strains, to which *C. elegans* is less susceptible.^48,49^ We also observed consistently smaller brood sizes for the interspecies (*C. elegans* × *C. brenneri*) cross compared to intraspecies matings, especially on the second day post-mating, (**Fig. 1B**).

**Figure 1.**
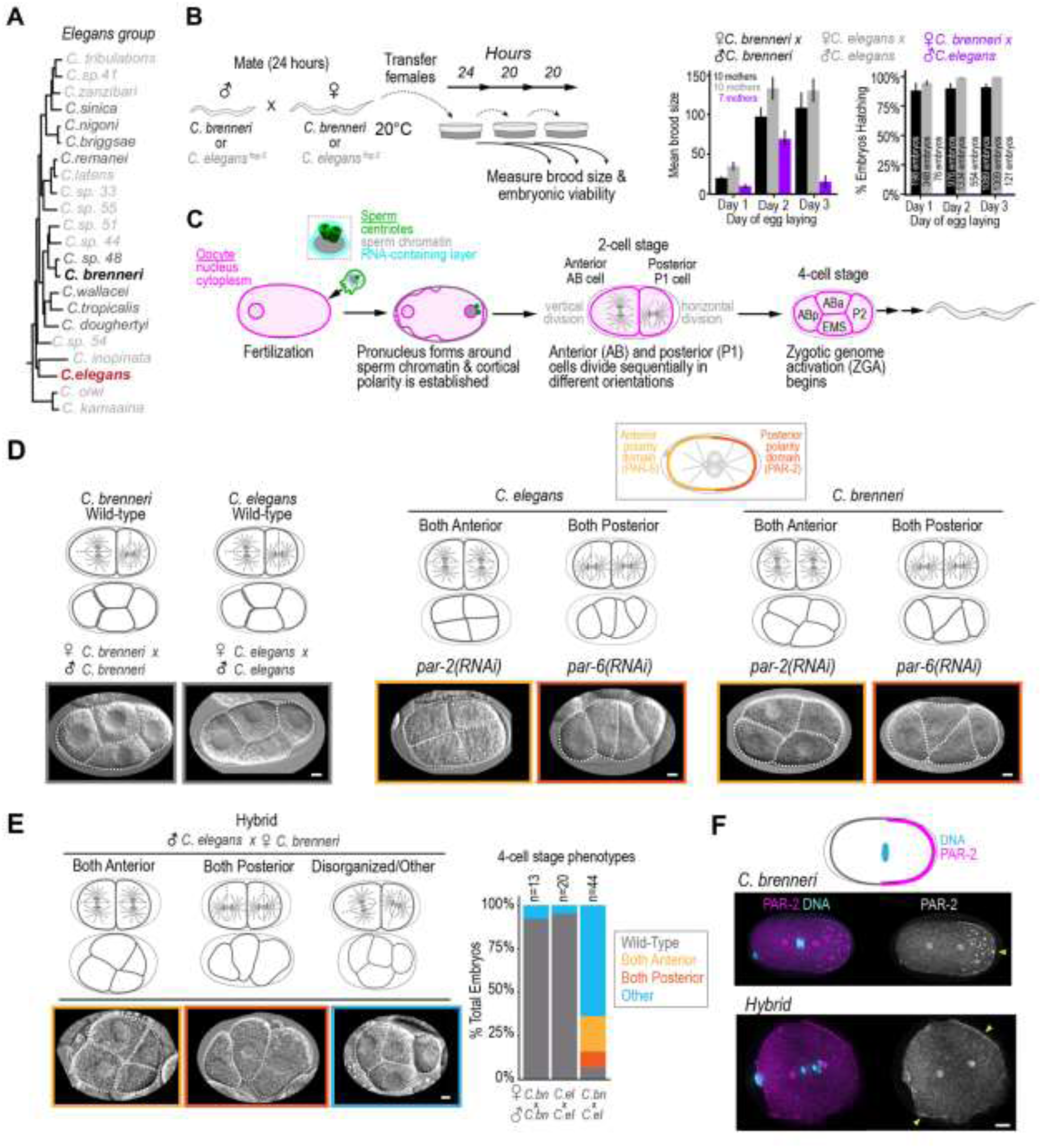
Hybrid embryos resulting from fertilization of *C. brenneri* oocytes with *C. elegans* sperm exhibit polarity defects prior to zygotic genome activation. (**A**) Phylogeny of the *Elegans* group of *Caenorhabditis*.^71^ *C. brenneri* females have been shown to be cross-fertile with *C. elegans* (*red*) males and males from the species marked in dark grey. (**B**) Schematic illustrating how embryos were collected after mating. Graphs show the average brood size and % hatching for the embryos collected during each day of egg laying. Day 1 is 0-24 hours post-mating; day 2 is 24-44 hours post-mating; day 3 is 44-64 hours post-mating. n refers to the number of embryos counted per cross. Error bars are ± 1 SE. **(C)** Schematic of the events between fertilization and the 4-cell stage when Zygotic Genome Activation begins. The sperm brings the sperm chromatin, which is not encased in a nuclear envelope but is instead embedded in an RNA-containing layer with a pair of centrioles,^72^ into the oocyte. Once in the oocyte, a nuclear envelope assembles around the sperm chromatin to form the sperm-derived pronucleus and the centrioles form centrosomes. The larger, anterior AB cells divides first followed by the smaller, posterior P1 cell, leading to the canonical arrangement of cells at the 4-cell stage. ZGA begins, when the sperm genome is first expressed. Hybrid incompatibilities prior to this stage are due to incompatibilities between the components derived from the sperm and oocyte. (**D**) Differential interference contrast (DIC) images of 4-cell stage embryos for: (*left*) control intraspecies *C. brenneri* and *C. elegans* embryos; (*middle*) *par-2(RNAi)* and *par-6(RNAi) C. elegans* embryos; and (*right*) and *par-2(RNAi)* and *par-6(RNAi) C. brenneri* embryos. For phenotypic quantification see *Fig. S1D*. The schematic on top illustrates the anterior PAR-6 (*yellow*) and posterior PAR-2 (*orange*) domains at the 1-cell stage in *C. elegans*. Schematics above the images show the expected cell and spindle orientations for two-cell and four-cell stage embryos for each condition. (**E**) DIC images illustrating the three non-wild-type classes of 4-cell stage phenotypes observed for hybrid embryos resulting from mating *C. brenneri* females with *C. elegans* males (Both Anterior, Both Posterior, and Disorganized/Other). Schematics above the images illustrate the cell and spindle orientations at the two-cell and four-cell stages for each embryo. The graph quantifies the percentage of embryos exhibiting each phenotype. (**F**) (*top*) Schematic illustrating the localization of PAR-2 and DNA in a control embryo. (*left*) Representative immunofluorescence images of *C. brenneri* (top; n = 25) and hybrid embryos (bottom; n = 9) stained for PAR-2 (*magenta*) and DNA (*cyan*). (*right*) Grayscale PAR-2 images with yellow arrowheads marking the PAR-2 domains. Scale bars, 5μm.

Work in *C. elegans* and other *Caenorhabditis* species has shown that zygotic transcription begins at the 4-cell stage.^20–23^ This makes the 4-cell stage a useful developmental boundary (**Fig. 1C**): defects observed at or before this point are likely caused by incompatibilities between the physical components derived from the oocyte and sperm, while defects arising later may reflect mismatches in the timing and regulation of zygotic gene expression or incompatibilities between zygotically expressed gene products. To determine whether defects in hybrid embryos appear before the 4-cell stage, or first emerge as embryos begin tissue-specific gene expression, we mated unmarked *C. brenneri* or *C. elegans* (*fog-2*) females to *C. elegans* males carrying fluorescent reporters that label nuclei in tissues derived from the three germ layers endoderm (*green*), mesoderm (*yellow*), and ectoderm (*red*)^50,51^ (**Fig. S1A**). As a control, unmarked *C. brenneri* males were also mated to *C. brenneri* females. Because the fluorescent reporters are only expressed later in development, we used differential interference contrast (DIC) microscopy to film embryos from just after fertilization through the 4-cell stage^52^ (**Fig. 1D-E**). Embryos were then incubated overnight, and a fluorescence z-stack was collected the following day to determine the point of arrest and whether the tissue-specific markers had turned on (**Fig. S1A-C**). Since we observed significant defects by the 4-cell stage, we begin by describing the results of our DIC analysis of early development.

DIC filming of early embryogenesis (schematized in **Fig. 1C**) revealed significant defects in hybrid embryos. To interpret these defects, we draw on prior analyses in *C. elegans*, where—as in most metazoans—oocytes lack centrioles and centrosomes,^24,25^ and the sperm contributes a pair of centrioles along with the paternal chromatin. After fertilization, these centrioles mature into centrosomes that remain associated with the sperm-derived pronucleus. The sperm pronucleus holds the centrosomes against the cell cortex, which allows them to deliver a signal that marks the embryo posterior and triggers the formation of two opposing cortical domains: a posterior domain containing a PAR (“Partitioning-defective”) protein complex that includes PAR-2, and an anterior domain containing a complex that includes PAR-6 (**Fig. 1D**).^33,34,53–55^ The first mitotic spindle forms along this axis and divides the embryo into a larger anterior (AB) cell and smaller posterior (P1) cell. During the second division, the AB spindle aligns vertically (perpendicular to the anterior-posterior axis of the embryo), while the P1 spindle aligns horizontally (parallel to the axis). This results in the stereotypical 4-cell arrangement observed in *Caenorhabditis* embryos (ABa on top of EMS in the middle, flanked by ABp and P2 on the anterior and posterior sides; **Fig. 1C,D**). Proper organization at this stage is essential for correct cell-cell signaling and fate specification.^34^

In *C. elegans*, knocking down *par-2* prevents formation of a functional posterior domain. As a result, the mitotic spindles in both daughter cells orient vertically—similar to the anterior AB cell— producing a 4-cell arrangement that resembles a 4-leaf clover (Both Anterior; **Fig.1D**, **Fig. S1D**).^34,53,54,56,57^ Conversely, knocking down *par-6* disrupts formation of the anterior domain, causing both spindles to orient horizontally—like the posterior P1 cell—resulting in a linear chain of cells (Both Posterior; **Fig. 1D**, **Fig. S1D**).^53,54,56^ Similar polarity defects were observed in *C. brenneri* embryos following RNAi of *par-2* and *par-6*, although the *par-6* knockdown produced Both Anterior and Both Posterior phenotypes in roughly equal numbers (**Fig. 1D**, **Fig. S1D**; **Video S1**). Polarity was intact in 92% (12/13) of *C. brenneri* and 95% (19/20) of *C. elegans* intraspecies embryos, whereas only 7% (3/44) of hybrid embryos from crosses between *C. elegans* males and *C. brenneri* females exhibited normal 4-cell stage organization. Instead, 21% (9/44) showed a Both Anterior phenotype, 9% (4/44) showed a Both Posterior phenotype, and 64% (28/44) exhibited aberrant arrangements that typically included an abnormally positioned P1 cell (Other; **Fig. 1E**). To more directly probe polarity establishment, we used a *C. elegans* anti-PAR-2 antibody^58^ that also labels the posterior cortex in *C. brenneri* 1-cell stage embryos (25/25). In hybrids, PAR-2 localization was abnormal: 3/9 embryos lacked a defined PAR-2 domain, and 5/9 showed incomplete or misplaced domains (**Fig. 1F**). Only one hybrid established a proper PAR-2 domain.

To determine whether polarity defects affected progression to mid-embryogenesis and zygotic expression of tissue-specific markers, we imaged hybrid embryos the following day using DIC and spinning disk confocal fluorescence microscopy (**Fig. S1A-C**). 12.5% (1/8) of hybrid embryos with a Both Anterior phenotype, 25% (1/4) with a Both Posterior phenotype, and 38% (10/26) with an Other aberrant 4-cell morphology phenotype turned on germ layer reporter expression prior to developmental arrest. In contrast, 100% (3/3) of hybrid embryos with normal 4-cell morphology expressed germ layer reporters before arrest (**Fig. S1A-C**).

We conclude that many hybrid embryos resulting from fertilization of *C. brenneri* oocytes by *C. elegans* sperm exhibit defects consistent with compromised anterior-posterior polarity by the 4-cell stage. These defects, which arise before zygotic transcription begins, suggest early incompatibility between sperm-and oocyte-derived components. Moreover, polarity defects compromise cell fate specification and greatly reduce the likelihood of activating tissue-specific gene expression.

### Delayed expansion of the sperm pronucleus leads to centrosome detachment and is the likely cause of polarity defects in *C. brenneri x C. elegans* hybrid embryos

Next, we wanted to understand the nature of the incompatibility that gives rise to the polarity defects in hybrid embryos. Cortical polarity is established at the one-cell stage when the sperm-provided centrioles recruit a pericentriolar material (PCM) scaffold that anchors γ-tubulin complexes for microtubule nucleation. When held against the cortex by the sperm pronucleus, the centrosomes provide a cue that establishes cortical polarity (**Fig. 2A**).^33,59–61^ To determine if *C. elegans* sperm centrioles can recruit PCM in *C. brenneri* oocytes, we used immunofluorescence to visualize γ-tubulin (which docks onto the PCM scaffold)^60,62^ and α-tubulin (to assess microtubule nucleation). In all embryos examined—*C. elegans* (12/12), *C. brenneri* (14/14), and hybrids (12/12)—centrosomes adjacent to the sperm pronucleus recruited γ-tubulin and nucleated α-tubulin-containing microtubules (**Fig. 2B**; **Fig. S2A**). Thus, *C. elegans* centrioles are competent to form functional centrosomes in *C. brenneri* oocytes. However, a key difference was observed in hybrids: while centrosomes remained attached to the sperm pronucleus in intraspecies embryos, one or both centrosomes were frequently observed detached in hybrids (11/12 embryos; **Fig. 2B**; **Fig. S2A**).

**Figure 2.**
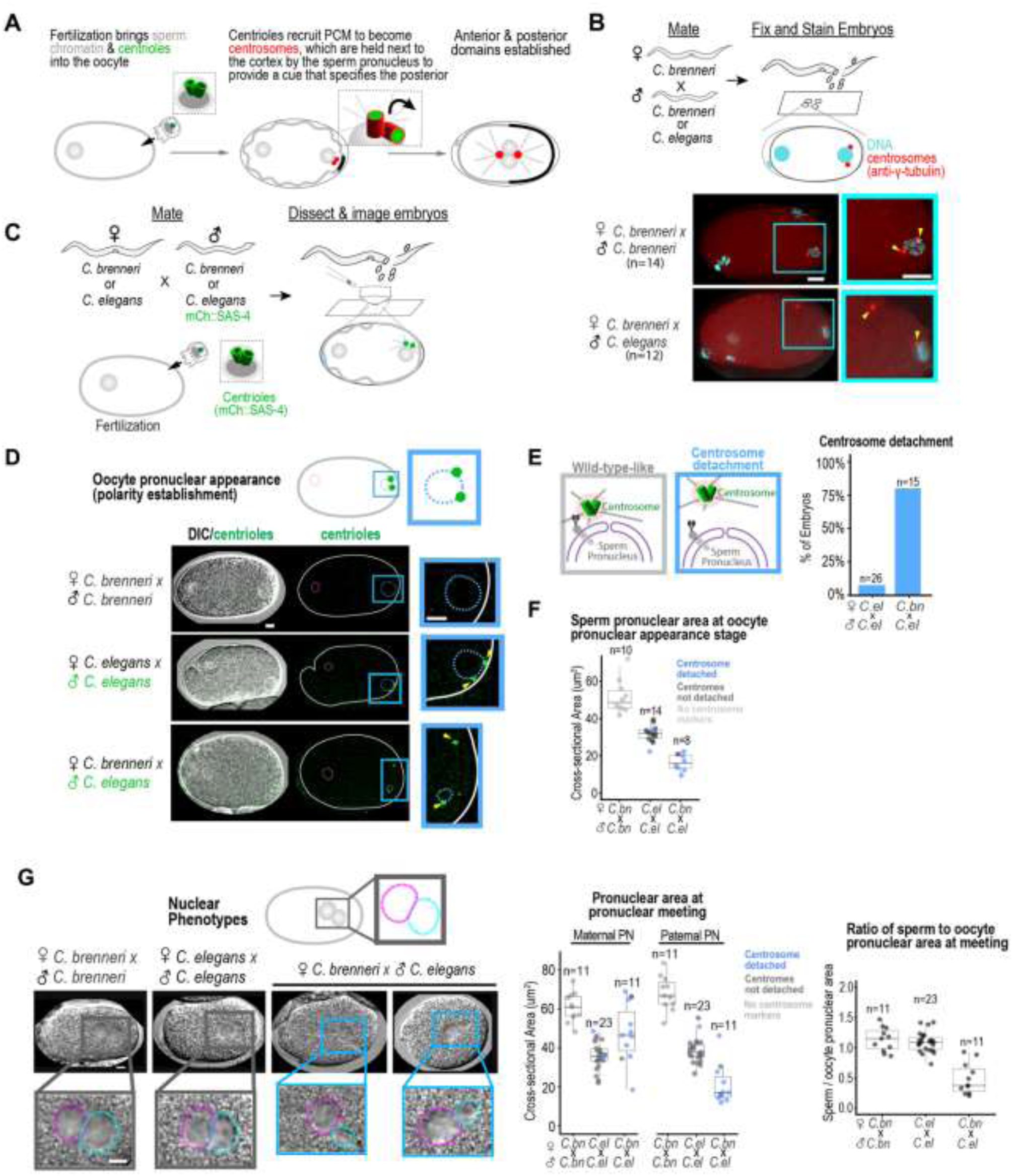
Sperm pronuclear expansion is delayed in hybrid embryos, leading to centrosome detachmet, which can cause polarity defects. (**A**) The schematic illustrates how fertilization triggers polarity establishment in the 1-cell embryo. The sperm brings in a pair of centrioles (*green*) that are converted to centrosomes via recruitment of a microtubule-nucleating scaffold (*red*) from the oocyte cytoplasm. The sperm pronucleus holds the centrosomes next to the cortex, where they provide a cue that specifies the embryo posterior. (**B**) (*top*) The schematic illustrates the mating regime and markers shown in the immunofluorescence images below. (*bottom*) Representative images of 1-cell stage *C. brenneri* (n = 14) and hybrid (n = 12) embryos after the conversion of centrioles to centrosomes, but prior to pronuclear migration, stained for DNA (cyan) and γ-tubulin (red). Insets are magnified 1.7X. γ-tubulin (red) is scaled equivalently in the two images. (**C**) The schematic illustrates the mating of *C. brenneri* or *C. elegans* females with *C. brenneri* males or with *C. elegans* males whose centrioles are stably marked with mCherry::SAS-4 (*green*) to allow monitoring of sperm centriole position throughout the first cell division. (**D**) Paired DIC/fluorescence overlay (*left*) and fluorescence-only (*right*) images of *C. brenneri*, *C. elegans,* and hybrid pronuclear appearance stage 1-cell embryos. The schematic on top illustrates the expected position of the mCherry::SAS-4 marked centrioles between the sperm pronucleus and cortex (*green*). The white solid lines trace the embryo. The white dashed lines trace pronuclei in DIC and fluorescence images. Insets are magnified 2.3X, and yellow arrowheads mark centrosomes. Image intensities for centrioles are scaled to highlight centriole position and cannot be compared across images. (**E**) The schematics depict wild-type-like attachment (*grey box*) and aberrant detachment (*blue box*) of centrosomes from the sperm pronucleus. The graph quantifies the percentage of embryos with centriole detachment (>9 μm separation) from the sperm pronucleus. (**F**) The graph plots the cross-sectional area of sperm pronuclei at the pronuclear appearance stage. The median (IQR) sperm pronuclear cross-sectional areas are 49 (39-58), 32 (28-37), and 16 (9-23) μm^2^ for *C. brenneri*, *C. elegans*, and hybrid embryos, respectively. Centriole detachment is indicated by blue circles, and wild-type-like centriole separation is indicated by dark gray circles. (**G**) DIC images of representative *C. brenneri*, *C. elegans*, and *hybrid* embryos at pronuclear meeting. Insets are magnified 2X (the cyan and magenta dotted lines outline the sperm and oocyte-derived pronuclei, respectively). The left graph plots the cross-sectional areas of oocyte and sperm pronuclei at pronuclear meeting for the indicated crosses. Centriole detachment is indicated by blue circles and wild-type-like centriole separation by dark gray circles. In *C. brenneri*, centrioles were not labeled, and detachment was not scored (*light gray circles*). Median (IQR) of female pronuclear cross-sectional areas are 61 (51-71), 36 (30-42), 46 (26-66) μm^2^ for *C. brenneri, C. elegans*, and hybrid embryos, respectively. Median (IQR) of male pronuclear cross-sectional areas are 67 (56-78), 38 (32-44), 17 (7-27) μm^2^ for *C. brenneri, C. elegans*, and hybrid embryos, respectively. Right graph plots the ratio of sperm to oocyte pronuclear area at pronuclear meeting. Median (IQR) ratios for sperm pronuclear area to oocyte pronuclear areas are 1.2 (0.9-1.5), 1.1 (0.9-1.3), 0.4 (0.01-0.73) for *C. brenneri*, *C. elegans*, and hybrid embryos, respectively. n refers to the number of embryos quantified. Scale bars, 5μm.

Since cortical polarity depends on centrosomes remaining near the cortex, we tracked centrosome behavior in living embryos. We mated unmarked *C. brenneri* or *C. elegans* (*fog-2*) females with C. elegans males expressing mCherry-tagged SAS-4 (a stably incorporated centriolar protein that does not turnover; **Fig. 2C**).^62^ Unmarked *C. brenneri* males were also mated to *C. brenneri* females as a control. During polarity establishment in *C. elegans* intraspecies embryos, the two mCherry::SAS-4 marked centrosomes separated as the sperm pronucleus came into proximity to the cortex (**Fig. 2D**). Both centrosomes remained attached to the sperm pronucleus during this process (Oocyte pronuclear appearance stage; 24/26 embryos; **Fig. 2D**; **Video S2**). Centrosome-pronuclear attachment is mediated by dynein anchored to the pronuclear envelope via the LINC complex,^63–65^ which pulls the centrosomes towards the pronucleus by walking along microtubules towards their minus ends (**Fig. 2E**). In contrast, in most hybrid embryos at this same crucial stage, one centrosome was already detached from the sperm pronucleus (>9 µm of separation) and migrated into the cytoplasm or along the cortex (12/15 embryos; **Fig. 2E**). Although they were typically recaptured prior to or during pronuclear meeting (**Video S2**), the centrosomes were often mispositioned relative to the two pronuclei at nuclear envelope breakdown (10/15 embryos; **Fig. S2B**). Nonetheless, both centrioles were successfully segregated to the daughter cells (**Fig. S2C**).

Transient centrosome detachment followed by recapture has been previously reported for perturbations that reduce pronuclear size.^52,66^ Experimental data support a model in which the sperm pronucleus has limited surface area for microtubule interactions, which limits attachment to a single centrosome until a size threshold is reached.^66^ To determine whether small sperm pronuclear size could be the cause of centrosome detachment in hybrids, we measured the cross-sectional area of the sperm and oocyte pronuclei at the time of pronuclear appearance and pronuclear meeting. At both stages, the cross-sectional area of the sperm pronuclei in hybrids was about half of that in intraspecies *C. elegans* embryos and of the corresponding oocyte pronuclei (**Fig. 2F,G**). We conclude that the *C. elegans* sperm-derived pronucleus is unable to properly expand in the *C. brenneri* oocyte cytoplasm, which leads to centrosome detachment and is the likely cause of the observed polarity defects.

### Hybrid embryos exhibit defects in polar body extrusion and spindle morphology

The above results indicate that centrosome detachment due to delayed sperm pronuclear expansion leads to mispositioned centrosomes at nuclear envelope breakdown (NEBD). Next, we tested if these effects cause defects in spindle orientation and morphology in hybrid embryos (**Fig. 3A**). To analyze spindle orientation, we measured the angle between the centrosome-to-centrosome axis of the spindle and the anterior-posterior axis of the embryo (**Fig. 3B**). In control embryos, 95% (161/168) of spindles were oriented within -7.5° to 10.6° of the anterior-posterior axis. In contrast, only 37% of spindles in hybrid embryos fell within this range. Hybrid embryos displayed a much wider orientation range, from -66° to 51° (**Fig. 3B**). These abnormal spindle angles are consistent with polarity defects causing large spindle oscillations during anaphase.^52,67,68^

**Figure 3.**
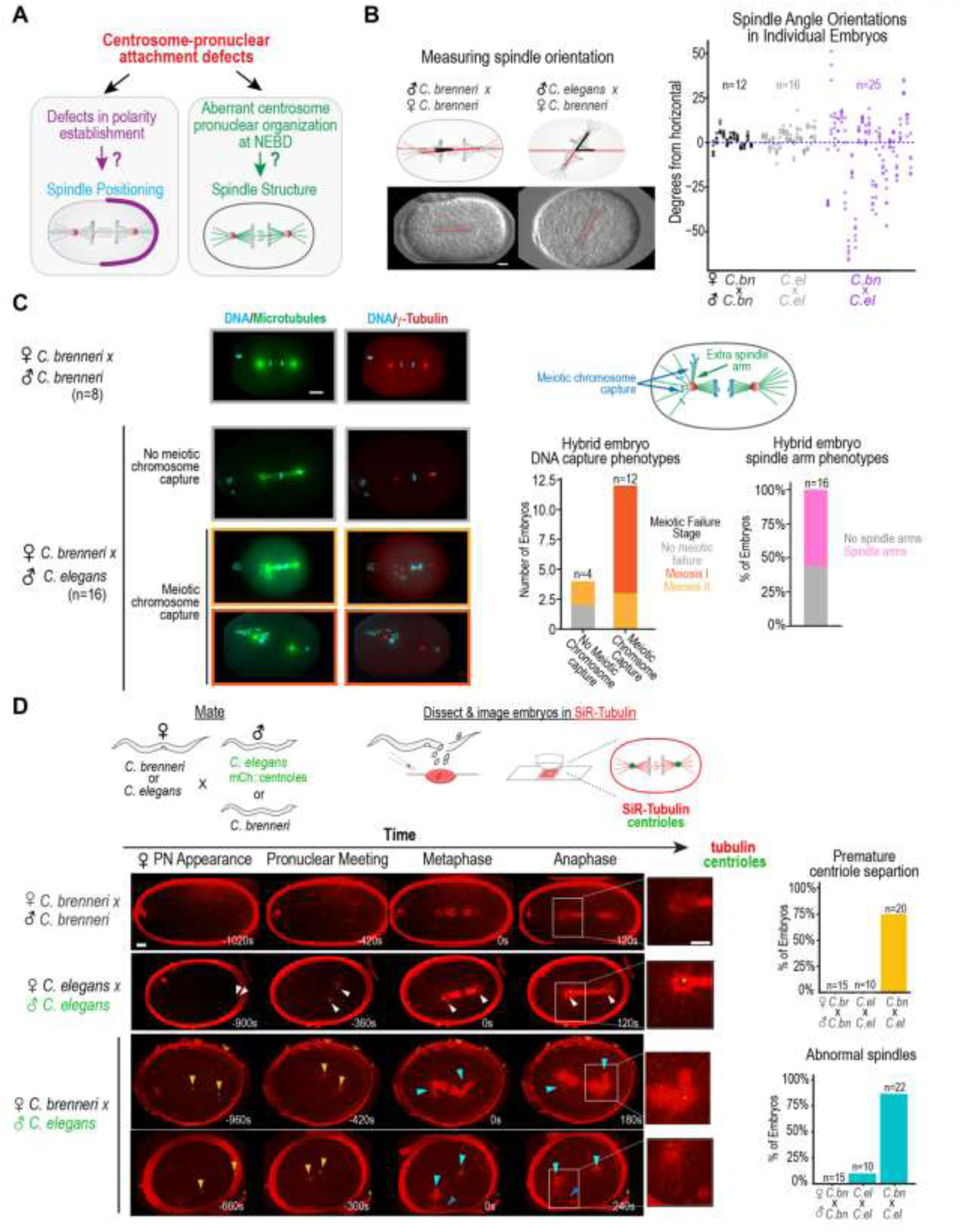
*Caenorhabditis* hybrid embryos exhibit defects in polar body extrusion and spindle morphology. (**A**) A schematic illustrating how detachment of centrosomes from the sperm pronucleus could contribute to defects in polarity establishment, spindle positioning, and spindle morphology in the early embryo. (**B**) Schematics (*top*) and images (*bottom*) illustrating how spindle orientation was assessed. At each timepoint, the angle between the centrosome-to-centrosome axis of the spindle (red dashed line) and the anterior-posterior axis of the embryo (red solid line) was measured. For each embryo, six measurements were made at 30-45 second intervals starting at metaphase. The graph plots the six measured angles for each embryo in a column. Angles were measured in *C. brenneri* (*black*), *C. elegans* (gray) or hybrid (*purple*) embryos. (**C**) Representative immunofluorescence images of *C. brenneri* (n = 8) and hybrid (n = 16) are shown to illustrate the range of hybrid phenotypes, which include abnormally structured spindles due to mispositioned centrosomes and spindles that have captured meiotic chromosomes due to failed polar body extrusion in meiosis I or meiosis II. The schematic (*top right*) illustrates the meiotic chromosome capture and ‘spindle arms’ phenotypes observed in embryos stained for DNA (*cyan*), microtubules (DM1-α) (*green*) and γ-tubulin (*red*). The graphs quantify the number of hybrids displaying each meiotic failure phenotype with/without meiotic chromosome capture (l*eft*) and the frequency of hybrid embryos exhibiting a ‘spindle-arm’ phenotype (*right*). (**D**) (*Top*) A schematic illustrating how *C. brenneri* or *C. elegans* females were mated with *C. elegans* males with mCherry::SAS-4-marked centrioles to enable live tracking of centrioles (*green*). As a control, *C. brenneri* females were also mated with *C. brenneri* males. Embryos were dissected into the vital dye SiR-Tubulin (*red*) to monitor microtubules. (*left*) Images are maximum intensity projections of representative intraspecies *C. brenneri* and *C. elegans* embryos and hybrids. Times are seconds relative to metaphase. Arrows indicate centrioles that separated with normal timing (white) or prematurely (yellow). Cyan arrows indicate abnormal spindle morphology. Dark blue arrows indicate meiotic chromosome capture. Insets show 2X magnifications of one anaphase spindle pole. Maximum intensity projections were made of all z-planes containing the centrosomes and spindle structures, and SiR-Tubulin intensities were scaled to best show spindle morphology and cannot be directly compared. (*right*) Graphs quantify the percentage of embryos exhibiting premature centriole separation (*yellow*), and abnormal spindle morphology (*cyan*) for the indicated conditions. Scale bar, 5μm.

To examine spindle morphology, we stained fixed embryos for γ-tubulin, microtubules, and DNA (**Fig. 3C**). Abnormal spindle structures—frequently including bundles of kinetochore microtubules extending out towards clusters of mispositioned chromosomes—were seen in 9/16 hybrid embryos, as might be anticipated given the centrosome detachment and positioning defects we had observed (**Fig. 2D**, **Fig. S2B**). Unexpectedly, 14/16 hybrid embryos displayed chromosomes with a meiotic-like morphology clustered near the centrosomal asters (**Fig. 3C**). In 9 of these embryos, no polar bodies were present, suggesting that polar body extrusion failed during both meiotic divisions. Five embryos had a single polar body, suggesting failure to extrude the second polar body (**Fig. 3C**). Thus, in addition to problems with pronuclear assembly and expansion, hybrid embryos resulting from fertilization of *C. brenneri* oocytes with *C. elegans* sperm frequently failed to extrude one or both polar bodies.

To visualize spindle dynamics in hybrid embryos in real time, we mated males from a *C. elegans* strain expressing the mCherry-tagged SAS-4 to unmarked *C. brenneri* or *C. elegans* (*fog-2*) females and dissected the worms into media containing SiR-Tubulin, a vital dye that stains microtubules. In control *C. elegans* and *C. brenneri* embryos, we observed well-formed centrosomal microtubule asters at both poles and robust kinetochore microtubule arrays aligned toward the centrally positioned chromosomes (**Fig. 3D**, **Video S2**). In contrast, 19/22 hybrid embryos displayed abnormal spindle morphologies. Many exhibited bundled microtubule arrays—or “spindle arms”—pointing in random directions, likely because centrosomes were attempting to capture clusters of mitotic or meiotic chromosomes that were in an atypical position when nuclear envelope breakdown occurred (**Fig. 3D**; **Fig. S2D; Video S2**). In 8/22 of hybrid embryos, the mitotic spindle microtubules appeared to engage remnants of the meiotic spindle near the embryo anterior and to them into the cytoplasm (**Fig. 3D**). The persistence of meiotic spindle remnants in the mitotic cytoplasm further supports the conclusion from our fixed cell analysis that hybrid embryos frequently fail to expel one or more of their polar bodies.^69^

### Similar early defects are observed in hybrid embryos from crosses between *C. brenneri* females and males from three other Elegans group species

We show above that hybrids produced by fertilizing *C. brenneri* oocytes with *C. elegans* sperm exhibit delayed expansion of the sperm-derived pronucleus. We suspect that this is because the *C. brenneri* oocyte cytoplasm struggles to release *C. elegans* sperm chromatin from its initial quiescent state or to properly encase it in a nuclear envelope with embedded nuclear pores. Hybrid embryos also frequently fail to extrude polar bodies. Embryos experiencing these problems tend to suffer subsequent defects in cell polarity and spindle morphology. To assess whether these early defects are common in hybrids with *C. brenneri,* we took advantage of the fact that C*. brenneri* females can be fertilized by males of several other species in the *Elegans* group.^40,42^ Nearly all such hybrids die during embryogenesis, a common outcome across interspecies *Caenorhabditis* crosses.^40,42^ To determine whether the severity of the developmental defects increases with phylogenetic distance, we analyzed hybrid embryos from two additional crosses: *C. brenneri* females mated with *C. sp. 48* and *C. remanei*. *C. brenneri* and *C. elegans* are estimated to have diverged ∼200 million generations (∼35 million years) ago, although these estimates have wide uncertainty.^70^ *C. sp. 48* is a sister species of *C. brenneri*^71^ and is about one-third as divergent from *C. brenneri* at the amino acid level as *C. elegans*. *C. remanei,* while it has a more recent common ancestor with *C. brenneri* than *C. elegans* (**Fig. 4A**), shows ∼17% greater protein divergence than *C. elegans*. Crosses between *C. brenneri* females and *C. sp. 48* males had brood sizes comparable to intraspecies crosses, but all embryos died during embryogenesis (**Fig. S3A**). Crosses between *C. brenneri* females and *C. remanei* males resulted in reduced brood sizes and nearly complete embryonic lethality (885/887 embryos died; **Fig. S3A**).

**Figure 4.**
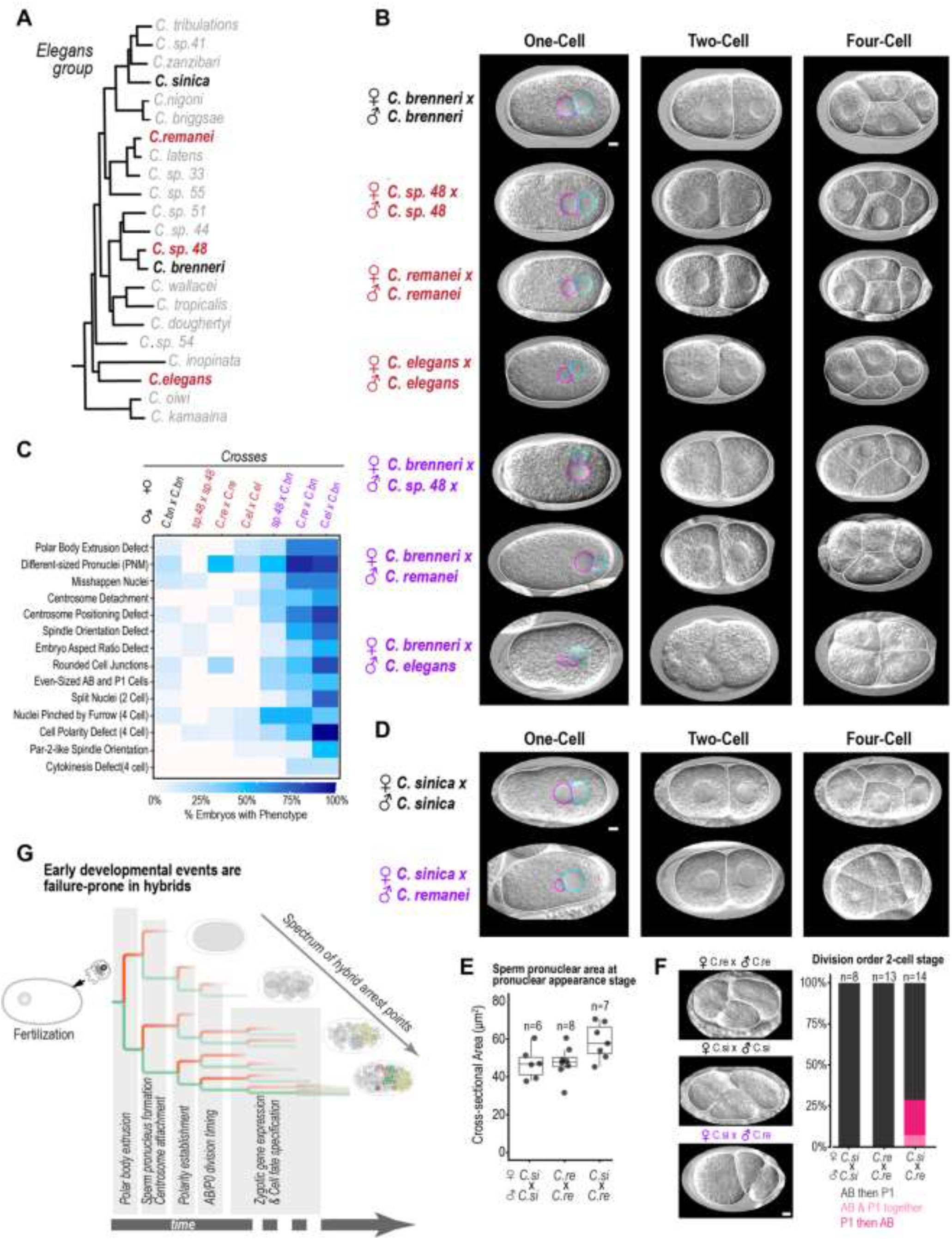
Similar phenotypic defects are observed in hybrid embryos generated by fertilization of *C. brenneri* oocytes with sperm from three Elegans group species. (A) (*Left*) Phylogeny of the *Elegans* group. In the experiments in *B* and *C*, *C. brenneri* (*black*) females were mated with males from three species: *C. remanei, C. sp. 48,* and *C. elegans (indicated in red). For D-F, C. sinica* (black) females were mated with *C. remanei* males. (**B**) DIC images from timelapse series of representative embryos showing them at the 1, 2, and 4-cell stages for all intraspecies (*red and black labels*) and interspecies crosses (*purple labels*). Solid white lines trace the outline of each cell; dotted cyan and magenta lines trace the sperm and oocyte-derived pronuclei, respectively. (**C**) The heatmap summarizes embryonic defects observed through the 4-cell stage for the indicated crosses; embryos were scored blinded to cross. Shading from white to dark blue indicates phenotype penetrance. Centrosome positioning defects are likely under-counted (compared to Figure 2D,*E*) because it is difficult to accurately follow centrosomes without a fluorescent marker. Aspect ratio scored as defective if it was outside of 2 standard deviations of the average across all wild-type embryos. (**D**) DIC images from timelapse series of representative embryos showing them at the 1, 2, and 4-cell stages for intraspecies *C. sinica* embryos (*C. remanei* shown in *B*) and the interspecies cross of *C. sinica* females to *C. remanei* males to (*purple labels*). Solid white lines trace the outline of each cell, dotted cyan and magenta lines trace the sperm and oocyte-derived pronuclei, respectively. The *C. remanei* sperm-derived pronucleus expands prematurely after it enters the *C. sinica* oocyte (see also *Fig. S3C*). (**E**) The graph quantifies the cross-sectional area of sperm pronuclei at the pronuclear appearance stage. Median (IQR) for sperm pronuclear cross-sectional areas are 46 (37-55), 47 (42-52), and 57 (43-71) μm^2^ for *C. sinica, C. remanei*, and hybrid embryos, respectively. (**F**) (*left*) After polarity establishment, the anterior AB cell divides prior to the posterior P1 cell, as observed in the dividing 2-cell intraspecies *C. remanei* and *C. sinica* embryos shown. In contrast, in the hybrid embryo shown, the P1 cell is undergoing cytokinesis before the AB cell. (*right*) The graph quantifies the AB/P1 division order in the embryos filmed for each condition. (**G**) A schematic illustrating the cascade of stochastic developmental failures in hybrid embryos. In hybrid embryos, a number of early embryonic events become more prone to failure, including polar body extrusion and the formation and expansion of the sperm pronucleus. These defects can, in turn, lead to centrosome detachment and defects in polarity establishment or 2-cell stage division timing. If hybrid embryos successfully navigate these early events, additional defects arise after the zygotic genome is activated and cell fate specification begins. Hybrid embryos exhibit a broad range of arrest points, and few survive to hatching. Scale bars, 5μm.

To characterize early developmental defects, we scored DIC movies of intraspecies embryos (*C. brenneri, C. sp. 48, C. remanei*, and *C. elegans*) as well as hybrid embryos resulting from fertilization of *C. brenneri* oocytes with sperm from the three other species for the presence of 20 distinct early embryonic defects (∼120 total movies; **Fig. 4C**; **Fig. S3A,B; Table S1**). A similar spectrum of early defects was observed for the hybrid embryos from the three crosses, including small and/or misshapen sperm pronuclei, detached or mispositioned centrosomes, and polar body extrusion defects. We also observed signs of disrupted cell polarity, such as equally sized AB and P1 cells at the 2-cell stage and abnormal 4-cell stage arrangements (**Video S3**). *C. brenneri* x *C. sp. 48* hybrids were often able to overcome their initial defects to reach a superficially normal 4-cell stage (**Fig. 4B**). Defects in the more distant *C. remanei* and *C. elegans* crosses were more frequent and severe compared to the sister species cross, suggesting that while early developmental processes are destabilized in hybrids, the severity of these disruptions increases with evolutionary distance between the parental species.

To determine whether such early defects are specific to hybrids generated by fertilization of *C. brenneri* oocytes or occur more broadly across the *Elegans* group, we also analyzed intraspecies *C. sinica* embryos and hybrids generated by fertilizing *C. sinica* oocytes with *C. remanei* sperm (**Fig. 4D-F**; **Fig. S3C,D**). Interestingly, while sperm pronuclei expanded slowly in hybrids generated by fertilizing *C. brenneri* oocytes with interspecies sperm (including *C. remanei*), *C. remanei* sperm pronuclei expanded prematurely after fertilizing *C. sinica* oocytes, reaching larger sizes than the sperm-derived pronuclei in either intraspecies cross (**Fig. 4D,E**; **Fig. S3C**). This observation indicates that sperm-derived pronuclear expansion can be either too slow or too fast in hybrids. Defects in centrosome positioning and spindle orientation were also observed in the *C. sinica x C. remanei* hybrids, which is consistent with the mismatch in sperm and oocyte pronuclear size (**Fig. S3D**). Moreover, in contrast to the normal pattern in *Elegans* group embryos—in which the larger anterior AB cell typically divides before the smaller posterior P1 cell— the posterior P1 cell divided first or simultaneously with the AB cell in ∼25% of the *sinica/remanei* hybrid embryos (**Fig. 4F**). Thus, both *C. brenneri* and *C. sinica* hybrids experience very early embryonic defects that eventually cascade into developmental failure.

## Conclusion

Our findings suggest that incompatibility between interacting components from oocytes and sperm is likely to be a common mechanism underlying hybrid incompatibility among nematodes of the *Caenorhabditis Elegans* group (**Fig. 4G)**. Two key drivers of incompatibility appear to be: (1) the ability of oocytes to properly control the timing of sperm-derived pronuclear expansion, and (2) the ability of fertilization to facilitate polar body formation. Electron microscopy has shown that sperm chromatin, along with the centrioles, is nested within an electron-dense, RNA-containing halo^26,28,29,72^ instead of being encapsulated by a nuclear envelope. In *C. elegans*, the sperm chromatin and centrioles initially remain quiescent while the oocyte chromosomes complete their meiotic segregation. The centrioles are activated to become centrosomes, and the sperm chromatin becomes encased in a nuclear envelope and initiates pronuclear expansion.^30,31^ Although the processes that remove the RNA halo and assemble an import-competent nuclear envelope around the sperm chromatin are not well understood, our data suggest that this transition is a common source of interspecies incompatibility. The defect may be in releasing the sperm chromatin from the RNA-containing halo or in the subsequent assembly of a nuclear envelope and nuclear pore insertion around the sperm chromatin. Our finding that divergence can disrupt the ability of oocytes to properly handle the packaged sperm chromatin has some similarity to prior work in other animal clades showing that divergence in centromeric or heterochromatic regions and their regulatory proteins can lead to an inability of the oocyte cytoplasm to segregate sperm-derived chromosomes.^14–19^ More work on the molecular specifics of these defects in controlling sperm pronuclear expansion will be needed to determine how similar or different the mechanisms driving these incompatibilities might be. We also observed frequent failures in polar body extrusion in hybrid embryos. Although, the cause is unclear, SPE-11—a sperm-provided protein that localizes to the RNA-containing sperm halo—is essential for polar body formation in *C. elegans*,^69,73,74^ suggesting that incompatible sperm factors could contribute to this failure. If the delay in sperm pronuclear expansion is caused by an inability to control the timing of disassembly of the RNA halo, the two defects may have a common origin.

To identify candidate proteins whose divergence may contribute to hybrid incompatibility, we extracted lists of genes in classes related to the phenotypes observed in hybrid embryos from Phenobank, a database that includes phenotypic data associated with *C. elegans* genes required for the first two embryonic divisions. We measured protein sequence divergence of these genes between *C. brenneri* and the other species in this study (**Fig. S4**). Low divergence proteins clustered with conserved proteins like the ubiquitin gene UBQ-1,^75^ and high divergence proteins clustered with sperm proteins like SPE-9 and SPE-11. Several proteins involved in the formation and structure of nuclear pore complexes, including MEL-28, NPP-7, NPP-4, NPP-23, and NPP-11,^76–82^ clustered with proteins exhibiting high divergence, raising the possibility that evolution at the amino acid level could have altered the association of one or more of these components with sperm chromatin in hybrid embryos and changed the timing of sperm pronuclear expansion.

Despite the fact that hybrid embryos suffer immediate problems right after fertilization, they do not die until later in development. The spectrum of varied developmental arrest points observed within each cross suggests that each problem incrementally destabilizes development and makes later events less robust and more prone to failure. Although development can often recover from moderate disruptions to its normal course,^83^ these hybrids do not have such resilience. While a few hybrid embryos may find a narrow pathway to hatching, most fall victim to a series of stochastic failures that build on each other, resulting in a spectrum of fatal outcomes (**Fig. 4G**).

## Supporting information

Key resources table

## ACKNOWLEDGEMENTS

We would like to thank members of the Rifkin, Oegema, and Desai labs for helpful discussions. We would like to thank Anthony Hyman for the generous gift of the PAR-2 antibody and Marie-Anne Félix for the generous gift of strain BRC20359 (*C. sp. 48*). We would also like to thank Anthony Ye and Aidan Linkins for their help. This work was supported by grants from the NIH (GM103782) and the NSF IOS (1936674) to S.A.R. and from the NIH to K.O. (GM147265). K.O. acknowledges salary support from the Ludwig Institute for Cancer Research. JB was partially supported by NIH/NIGMS (T32 GM127235). Some strains were provided by the CGC, which is funded by NIH Office of Research Infrastructure Programs (P40 OD010440).

## AUTHOR CONTRIBUTIONS

Conceptualization: J.B., R.G., K.O., S.A.R.; Methodology: J.B., R.G., K.O., S.A.R.; Formal analysis: J.B., R.G.; Investigation: J.B., R.G.; Resources: S.A.R., K.O.; Data curation: J.B., R.G.; Writing - original draft: J.B., R.G., K.O., S.A.R.; Writing - review & editing: J.B., R.G., A.D., K.O., S.A.R.; Visualization: J.B., R.G.; Supervision: R.G., S.A.R., K.O.; Project administration: S.A.R., K.O.; Funding acquisition: S.A.R., K.O.

## SUPPLEMENTAL FIGURE TITLES AND LEGENDS

**Figure S1.**
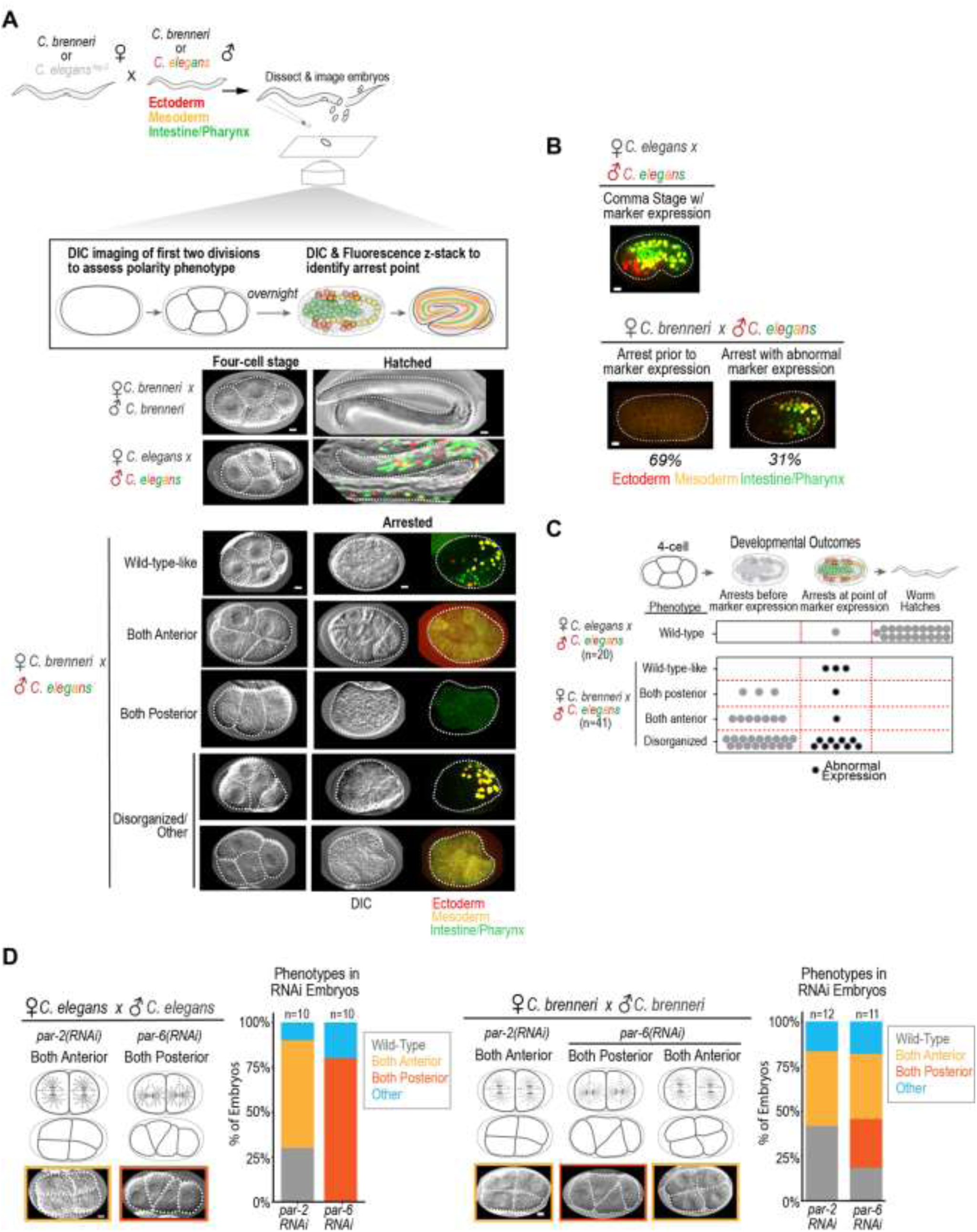
Polarity defects at the 4-cell stage correlate with the failure of hybrid embryos to reach mid-embryogenesis and turn on tissue-specific markers, related to Figure 1. **(A)** The schematic illustrates the experimental workflow to correlate early defects before ZGA, which occurs at the 4-cell stage, with the ability of embryos to reach mid-embryogenesis and turn on tissue-specific markers. Briefly, *C. brenneri* or *C. elegans* (*fog-2)* females were mated to males from either *a C. elegans* strain in which the genome encodes fluorescent reporters that mark nuclei in the endoderm (*green*), mesoderm (*yellow*), and ectoderm (*red*) or *C. brenneri (unmarked)*. DIC imaging was used to film embryos through the 4-cell stage. A spinning disk confocal fluorescence z-stack was collected of the same embryos 20-24 hours later to identify the point of arrest and determine whether the tissue-specific markers had turned on. Endpoint images for larval worms are scaled to best highlight marker expression, and intensity levels are not directly comparable to the arrested hybrid embryos. **(B)** Representative maximum intensity projections of spinning disc confocal z-stacks of hybrid embryos that arrested prior to (*bottom left*) and after (*bottom right*) the onset of marker expression compared to the comma stage in control *C. elegans* intraspecies embryos (*top*). Image projections are scaled to best highlight marker expression and intensities cannot be directly compared between *C. elegans* and hybrid embryos. (**C**) Summary table displaying the results of the phenotypic analysis for the *C. elegans* and *C. brenneri* x *C. elegans* hybrid embryos. While the majority of *C.* elegans embryos (19/20) showed a wild-type 4-cell phenotype and hatched, most hybrid embryos (38/41) showed a non-wild-type-like 4-cell phenotype, and none hatched. All 3 of the hybrid embryos that exhibited a wild-type-like 4-cell orientation survived to express markers, whereas embryos exhibiting more severe polarity phenotypes rarely made it to marker expression (Both Anterior (1/4); Both Posterior (1/9)). Hybrid embryos that fell into the Disorganized/Other category had an intermediate level of success with 9/26 surviving to the point of marker expression. **(D)** DIC images of representative 4-cell stage embryos are shown for *par-2(RNAi)* and *par-6(RNAi) C. elegans* (*left*) and *C. brenneri* (*right*) embryos. The schematics highlight the 2-cell division planes giving rise to the 4-cell phenotypes, and the graphs show the phenotype frequencies for each condition. Scale bars, 5μm.

**Figure S2.**
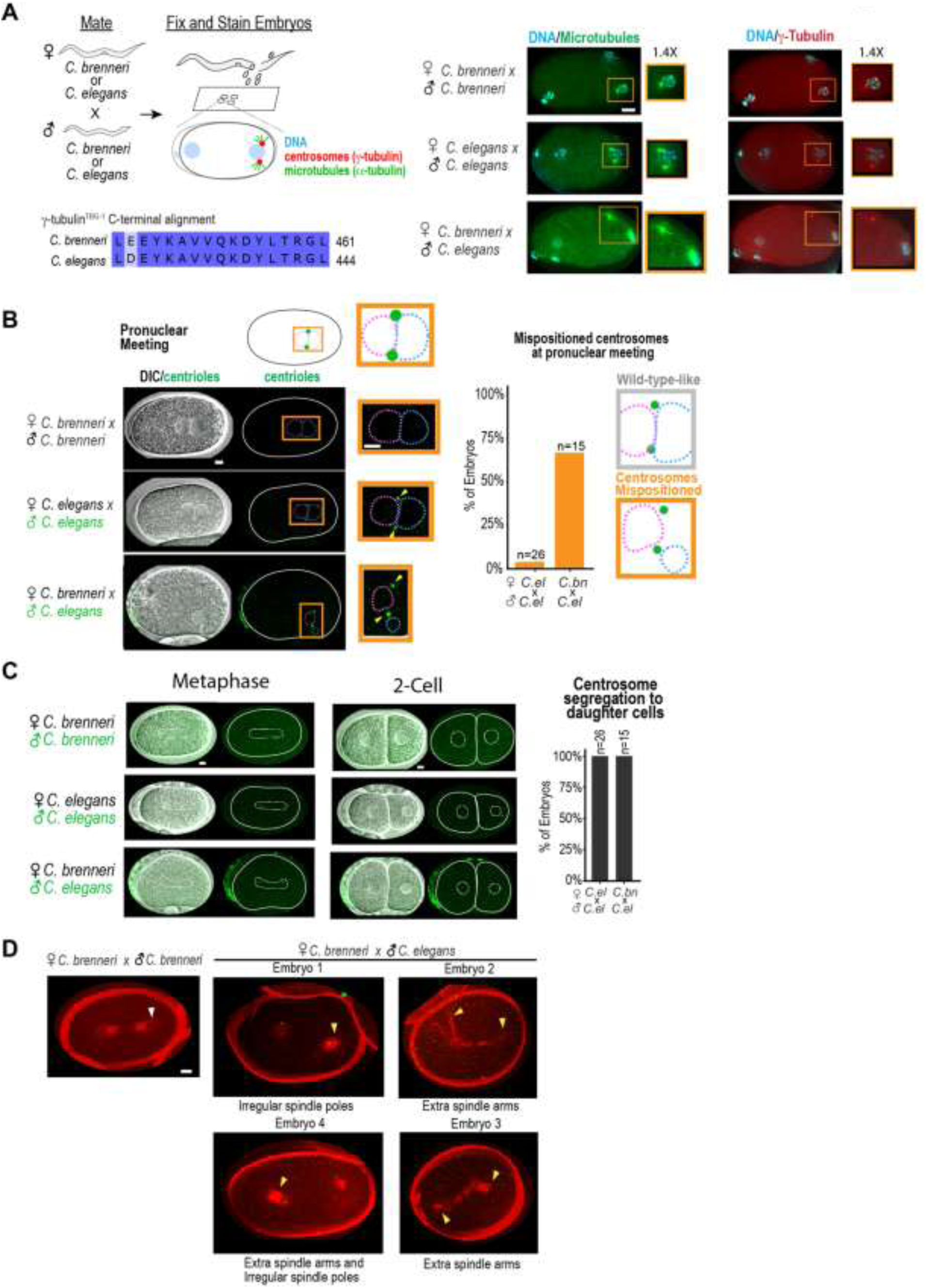
Hybrids exhibit defects in centrosome-pronuclear attachment and spindle morphology, related to Figures 2 and 3. **(A)** (*left*) The schematic illustrates the mating regime and the predicted staining patterns of α-tubulin, the centrosomal marker γ-tubulin, and DNA. An alignment of the C-terminal peptide of *C. elegans* γ-tubulin (TBG-1), against which the antibody was raised, to the equivalent region of the *C. brenneri* protein. (*right*) Representative images of 1-cell stage *C. brenneri* (n = 14), *C. elegans* (n = 12), and hybrid (n = 12) embryos after the conversion of centrioles to centrosomes but prior to pronuclear migration stained for DNA (*cyan*), α-tubulin (DM1α) (*green*), and γ-tubulin (*red*). Insets are magnified 1.4X. One or both centrosomes were usually detached in hybrids (11/12 embryos). Maximum intensity projections were scaled to best highlight protein localization, so intensity levels are not directly comparable across images. (**B**) (*left*) Paired DIC/fluorescence overlay and fluorescence-only images of *C. brenneri*, *C. elegans,* and hybrid pronuclear meeting stage 1-cell embryos. The schematic on top illustrates the expected position of the mCherry::SAS-4 marked centrioles (*green*). White solid lines trace the embryo. Dashed lines trace pronuclei; in fluorescence images, the sperm and oocyte-derived pronuclei are traced in blue and magenta, respectively. Insets are magnified 2.3X. Yellow arrowheads mark centrosomes. Image intensities are scaled to highlight centriole localization and cannot be compared across images. (*right*) The graph quantifies the percentage of embryos with wild-type-like (*gray box*) and mispositioned (*orange box*) centrosomes at pronuclear meeting. **(C)** Paired DIC/fluorescence overlay (*left*) and fluorescence-only (*right*) images of *C. brenneri*, *C. elegans,* and hybrid embryos at metaphase of the first division and the 2-cell stage. White solid lines outline the embryos and white dotted lines outline the spindle (metaphase) or nuclei (2-cell). Images are different timepoints taken from timelapse series of the same embryos shown in Figure 2D, *Figure S2B*. Image intensities for centrioles were scaled to best show centrosome localization and cannot be directly compared across embryos. The graph plots the percent of embryos that show the correct segregation of one sperm-derived centriole into each daughter cell (26/26 embryos in *C. elegans*; 15/15 embryos in hybrids). (**D**) *C. brenneri* females were mated with *C. brenneri* males as a control or with *C. elegans* males with mCherry::SAS-4-marked centrioles (*green*) and were dissected into the vital dye SiR-Tubulin (*red*) to monitor microtubules as described in Figure 3D. Images are maximum intensity projections that capture SiR-Tubulin-stained spindles (*red*) in a *C. brenneri* embryo (*top, left)* and four examples of hybrid embryos at anaphase. The white arrowhead indicates normal spindle morphology, whereas yellow arrowheads point to spindle poles or extra spindle arms in hybrid embryos. Embryos categorized as having abnormal spindles in the graph in Figure 3D displayed phenotypes similar to the ones shown here. Intensities were scaled to best show SiR-Tubulin signal and cannot be compared across embryos. Centriole intensities are scaled the same across images. *C. brenneri* embryos do not contain marked centrioles. Scale bars, 5μm.

**Figure S3.**
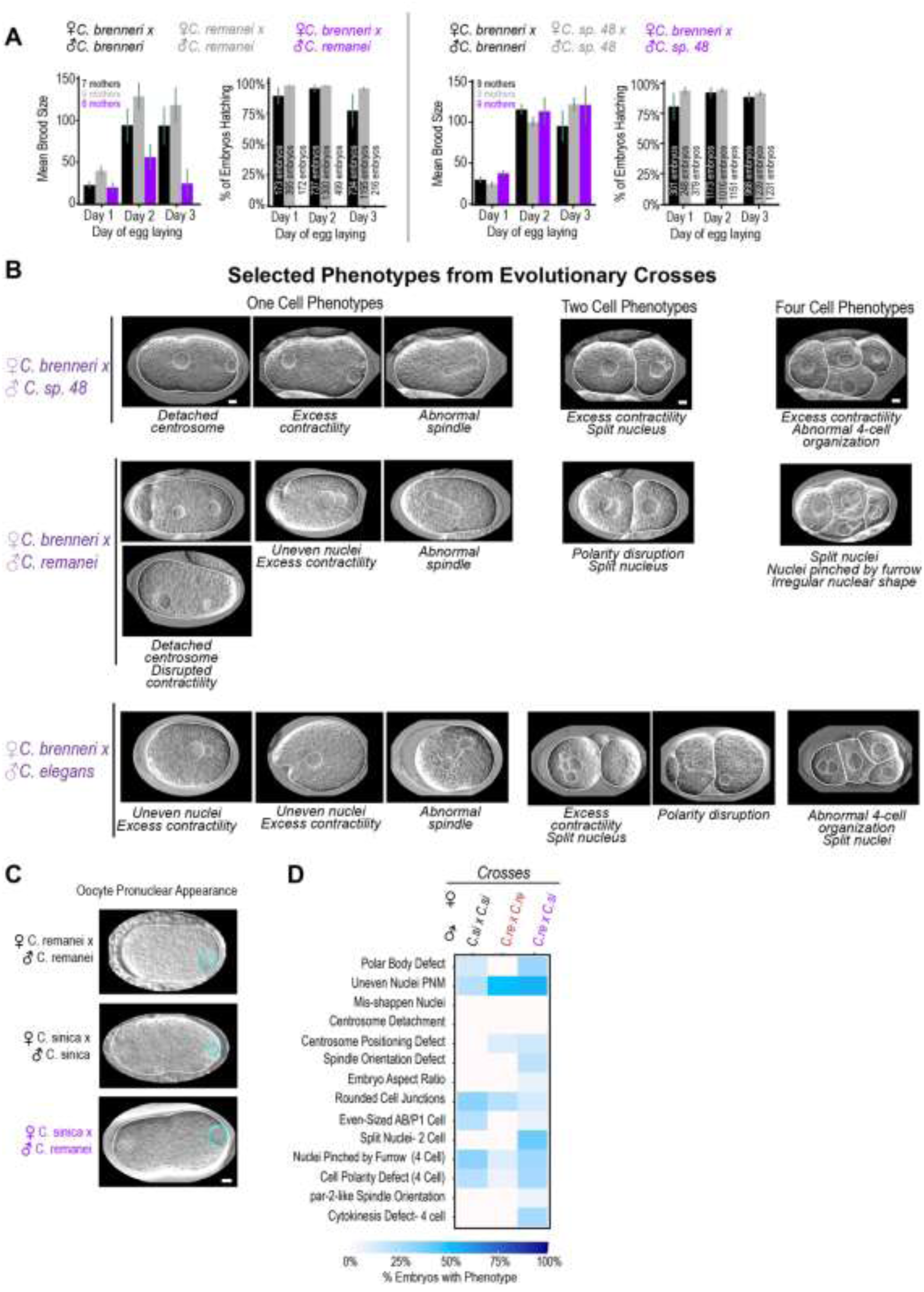
Gallery of hybrid phenotypes, related to Figure 4. (**A**) Graphs show the average brood size and % hatching for the embryos collected during the indicated days for the indicated species (*grey*, *black*) or hybrid (*purple*) cross. Day 1 is 0-24 hours post-mating; day 2 is 24-44 hours post-mating; day 3 is 44-64 hours post-mating. n refers to the number of embryos counted per cross. Error bars are ± SE. (**B**) The gallery highlights abnormal phenotypes found in 1, 2, and 4-cell hybrid embryos from *C. brenneri* females crossed with males from *C. sp. 48* (*top*), *C. remanei* (*middle*), and *C. elegans* (*fog-2(q71)*; *bottom*). White solid lines outline the embryos and white dotted lines outline the pronuclei, spindle, and nuclei. Yellow arrowheads highlight detached centrosomes. (**C**) DIC images of embryos at the oocyte pronuclear appearance stage for the indicated crosses. Solid white lines trace the perimeter of the embryos and dashed cyan dashed lines trace the perimeter of the sperm-derived pronuclei. (**D**) The heatmap summarizes embryonic defects observed through the 4-cell stage for the indicated crosses; embryos were scored blinded to cross. Shading from white to dark blue indicates phenotype penetrance. Scale bars, 5μm.

**Figure S4.**
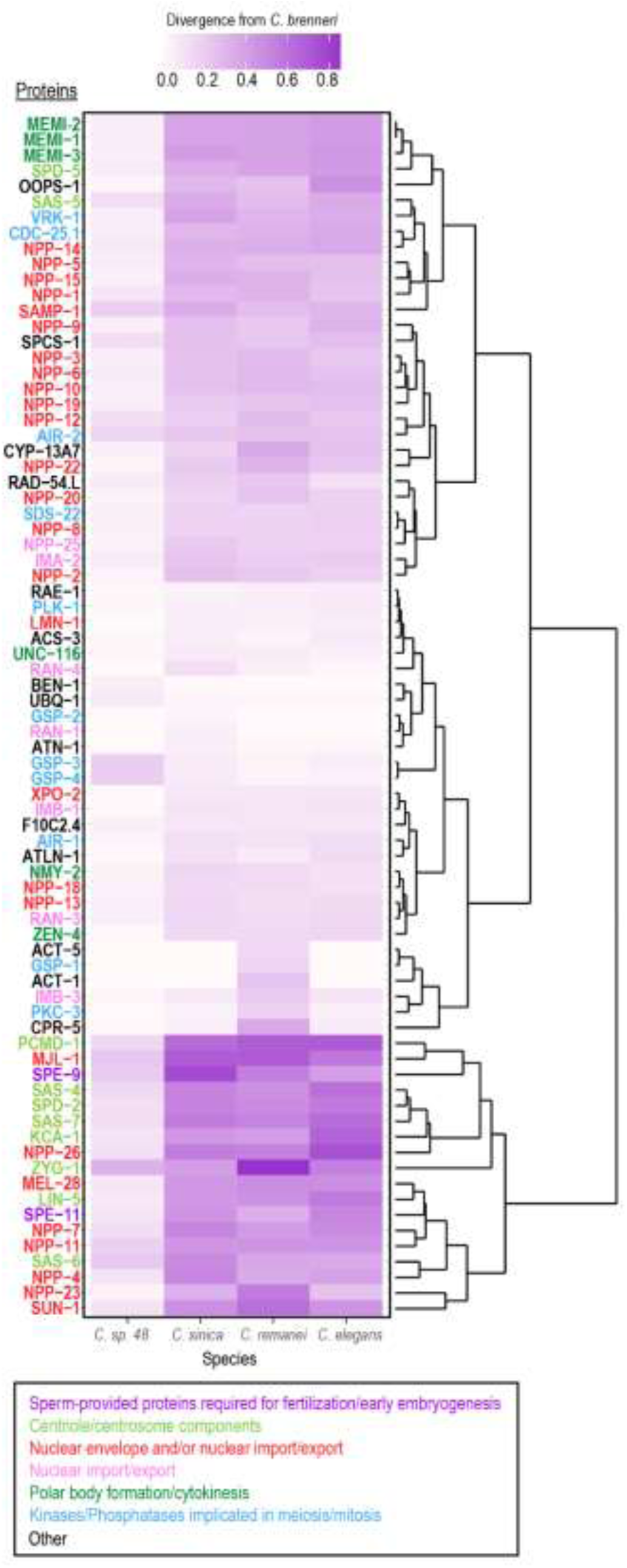
Sequence divergence in early embryonic proteins, related to Figure 4. (**A**) Heat map of protein sequence divergence scores of early embryonic proteins between *C. brenneri* and *C. sp. 48*, *C. sinica*, *C. remanei*, and *C. elegans*. Scores represent level of divergence from *C. brenneri* protein sequences, while the phylogeny clusters protein divergence scores based on similarity across all species compared to *C. brenneri*. The level of divergence is 1 – BLOSUM62 score (Biopython^84^). Gene names are color-coded as indicated in the key below the heat map. Proteins encoded by genes in the Phenobank phenotypic classes pronuclear appearance, asymmetry of division, centrosome attachment, and spindle assembly were subjected to pairwise protein sequence divergence analysis. To compare protein sequence divergence between *C. brenneri* and the species crossed with it in this study, as well *C. sinica*, we pulled out and clustered *C. brenneri* protein pairwise divergence scores versus all other species. Protein sequences clustered in 4 groups, with low divergence genes clustering with conserved genes like UBQ-1,^75^ and high divergence genes clustering with sperm genes like SPE-9 and SPE-11. Interestingly, a number of proteins involved in the formation and structure of nuclear pore complexes, such as MEL-28, NPP-7, NPP-4, NPP-23, and NPP-11^76–82^ clustered with high divergence genes. We note that SPE-11, a sperm-provided protein implicated in polar body formation^69,74^ and KCA-1, a kinesin-1 binding protein implicated in maintaining sperm centrioles in a quiescent state until after the second polar body is extruded during anaphase of meiosis II,^31^ also exhibit high divergence. While this protein sequence divergence analysis is biased towards genes associated with phenotypes we see in hybrids, it represents a starting point to identify proteins that may play a functional role in hybrid incompatibility between *C. brenneri* females and males of other species in the *Elegans* group.

**Video S1. Similar phenotypes are observed after RNAi of *par-2* and *par-6* in *C. elegans* and *C. brenneri* (related to Figure 1D**, **Figure S1D).** Timelapse sequences of *C. brenneri* (*top row*) and *C. elegans* embryos (*bottom row*). Control embryos are shown on the left, *par-2(RNAi)* embryos in the middle and *par-6(RNAi)* embryos on the right. Still images of the *C. brenneri par-6(RNAi)* and *par-2(RNAi)* embryos shown here are also shown in Figure 1D. Cleavage sites in the *par-2(RNAi)* (*yellow arrows*) and *par-6(RNAi)* (*orange arrows*) embryos are indicated. 26 x 1μm z-stacks were collected every 30 seconds and movies were created by compiling the best z-slice from the z-stack collected at each timepoint. Playback frame rate is 5 frames/second.

**Video S2. The sperm pronucleus remains small leading to centrosome detachment in hybrids of *C. brenneri* females and *C. elegans* males (related to Figure 2C-E and Figure S2B,C)** (00:00 – 00:21 seconds). Representative timelapse sequences showing centriole position and nuclear size during the first cell division in a *C. elegans* embryo (*top row*) and a *C. brenneri* x *C. elegans* hybrid (*bottom row*). Still images of the wild-type and hybrid embryos shown are also included in Figure 2D and *Figure S2B*. Yellow arrows point to the centrosomes. Black dotted lines trace the outline of the pronuclei. Movies are composed of a single z DIC slice overlaid with fluorescence maximum intensity projections of 19 x 1.5 μm z-stacks acquired every 30 seconds. Intensity values for centrioles were scaled to best highlight centrosome positioning are not comparable between different embryos. Playback rate is 5 frames/s. Aberrant spindle morphologies are observed in hybrid embryos of *C. brenneri* females and *C. elegans* males (related to Figure 3D and Figure S2E) (00:22 – 00:32 seconds). Representative timelapse sequences showing centrioles (*green*) and microtubules (*red*) in reference *C. brenneri* (first column) and *C. elegans* (second column) embryos along with two *C. brenneri* x *C. elegans* hybrids. Sequences run from ooctye pronuclear appearance through the first cell division. Still images of the embryos shown are also included in Figure 3D. The centrioles (*yellow arrows*), aberrant spindle morphology (*cyan arrows*), and meiotic spindle capture (*dark blue arrows*) are indicated. A 27 x 1 µm z-stack was collected every minute, and movies are composed of maximum intensity projections of the subset of z-planes that best show centrosomes and spindles for each timepoint. Intensity values for spindles are not comparable between different embryos because of the variable amount of dye that may enter the embryo. Playback speed is 5 frames/second.

**Video S3. A similar suite of early defects is observed in hybrids between *C. brenneri* females and males from three Elegans group species (related to Figure 4B-D**, **Figure 1D,E**, **Figure S1A and Figure S3B)** (00:00 – 00:30 seconds). Representative timelapse sequences of the early embryonic cell divisions in wild-type *C. brenneri*, *C. sp. 48*, *C. remanei* and *C. elegans* embryos (top row) along with their hybrids (*bottom row*). Still images of embryos shown are also included in Figure 4B, *Figure S1A* (*Disorganized/other)* and *Figure S3*. Black dotted lines trace pronuclei. Movies are composed of single z-slices chosen from 26 x 1 μm z-stacks acquired every 30sec. Playback rate is 5 frames/second. *C. sinica* hybrids are characterized by polarity and cell division timing defects (related to Figure 4D-F and Figure S3C,D) (00:31 – 00:55 seconds). Representative timelapse sequences of the early embryonic cell divisions in wild-type *C. sinica* and *C. remanei* (top row) along with their hybrid (bottom row). Stil images of wild-type embryos shown are also included in Figure 4F. Black dotted lines trace pronuclei. Movies are composed of single z-slices chosen from 26 x 1 μm z-stacks acquired every 30sec. Playback rate is 5 frames/second.

**Table S1.** Description of manual scoring data and measured features to Figures 3 and 4, Figure S3 and STAR Methods. In the All Scored Embryos tab, each cross type is highlighted in a different color. In the Considered Phenotypes tab, rows highlighted in yellow consistently showed a phenotype during embryo scoring.

## STAR METHODS

### RESOURCE AVAILABILITY

#### Lead contact

Further information and requests for resources and reagents should be directed to and will be fulfilled by the lead contact, Scott Rifkin (^31,32^).

#### Materials availability

All strains and other reagents generated in this study are freely available from the lead contact upon request. *C. brenneri, C. remanei, C. sinica,* and *C. elegans* (*fog-2*) strains can be obtained from the Caenorhabditis Genetics Center (CGC). *C. sp. 48* was a gift from M.-A. Félix.

#### Data and Code Availability

- The custom computer code generated for this project is publicly available through https://gitlab.com/evodevosyslabpubs/Bloom_etal_2025 and will be deposited at Zenodo
- Any additional information required to reanalyze the data reported in this paper will be available from the lead contact upon request

## EXPERIMENTAL MODEL AND STUDY PARTICIPANT DETAILS

*C. elegans, C. brenneri, C. sp. 48, C. sinica,* and *C. remanei* strains were maintained at 20°C on standard Nematode Growth Media (NGM) plates seeded with OP50 bacteria. The genotypes of the *C. elegans* strains used in this study are described in Reagents and Resources.

## METHOD DETAILS

### Strains

The following strains were used for experiments (see key resources table for details): *C. brenneri* wild-type LKC28; *C. sp. 48* wild-type BRC20359; *C. remanei* wild-type EM464; *C. sinica* wild-type JU1201; and *C. elegans* JK574 females, which has a *fog-2* mutation that makes hermaphrodites females.

### Mating

For all crosses, L4-stage females were placed with males on a 35mm plate seeded with OP50 and left overnight at 20C for mating; females were dissected the following day. A 1:2 ratio of females to males was used for all crosses except for interspecies crosses with *C. elegans* males, for which a 1:3 ratio was used because C*. elegans* males are worse at mating than dioecious males.^85,86^

### Dissections

Gravid females were dissected in Boyd Buffer (59.9 mM NaCl_2_, 32.2 mM KCl, 2.8 mM Na_2_HPO4, 1.8 mM CaCl_2_, 5mM HEPES pH 7.2, 0.2% glucose, 2.1 mM MgCl_2_;)^57,87^ and were transferred by mouth pipette to a 2% agarose pad made in Boyd Buffer for imaging. An 18×18mm coverslip was placed over the pad and the edges of the coverslip were sealed with VALAP (1:1:1 Vaseline, Lanolin, and Parafin) to prevent drying out. We compared five different osmotic support buffers (meiosis media, Boyd buffer, 0.5X Egg Salts, 0.7X Egg Salts, and 1X Egg Salts)^57,87–89^ but no rescue of early arrest phenotypes was observed for any of them.

### Brood size & embryonic viability measurements

To measure the number of embryos laid and assess their viability (defined hatching), individual L4 females were placed on a 35 mm NGM plate seeded with OP50 along with 2 or 3 males (see Mating section) and left to mate overnight at 20°C. After 24 hours, the males were removed and the females were moved to a second 35mm plate. Females were transferred again to a third 35 mm plate 20 hours later (44 hours after the start of mating). After 64 hours, the females were removed from the third plate. After each transfer, the number of freshly laid embryos on the plate from which the female was removed was counted and the plate was returned to 20°C. Since viable embryos hatch within 20 hours of being laid we waited 20-24 hours after first count was made and then counted the number of hatched and unhatched embryos to measure viability.

### RNA production

The *C. brenneri* orthologs of the *C. elegans par-2* and *par-6* genes were identified based on annotation in WormBase; ortholog identity was confirmed using reciprocal BLAST. DNA templates for generating dsRNAs were generated by PCR using primers designed using Primer3 (https://primer3.ut.ee/) to amplify a 400-800 bp region of each gene from genomic DNA (see KEY RESOURES Table for sequences). Primers contained T3 or T7 promoters to enable transcription reactions. Primers for dsRNAs targeting the *C. elegans* genes were the same as those employed in a prior RNAi-based screen.^52^ PCR reactions were cleaned and used as templates in T3 and T7 transcription reactions. T3 and T7 RNA products were mixed at equimolar amounts, cleaned, and annealed by adding 3X Soaking buffer (32.7 mM Na_2_HPO_4_, 16.5 mM KH_2_PO_4_, 6.3 mM NaCl, 14.1 mM NH_4_Cl) to a final concentration of 1X and incubating reactions at 68C for 10 minutes then 37°C for 30 minutes.

### RNA interference

For *C. elegans*, larval (L4 stage) female (JK574) worms were injected with dsRNA in the body cavity and left to recover at 20°C for 4 hours before singling and mating with male *C. elegans* (JK574) worms at 20°C and left overnight before imaging. For *C. brenneri*, larval (L4 stage) females (LKC28) were mated to male *C. brenneri* (LKC28) worms on a 35mm OP50 plate overnight at 20°C. Gravid females were injected with dsRNA in both gonad arms, left to recover for 3 hours (20°C), before singling and leaving overnight at 20°C for 22 hours before imaging.

### Imaging and analysis of early and late embryogenesis

After mating, dissection, and mounting as described above, we monitored early embryogenesis using differential interference contrast (DIC) optics to acquire 26 x 1µm z-stacks at 30-45 second intervals. In most experiments, embryos were imaged through the four-cell stage. Images were acquired using either an inverted Zeiss Axio Observer Z1 system equipped with a Yokogawa CSU-X1 spinning-disk, 63X 1.40 NA Plan Apochromat lens (Zeiss), and a QuantEm: 512SC camera (Teledyne Photometrics), or on a Nikon Ti2 microscope equipped with a Yokogawa CSU-X1 spinning disk, a 60X, 1.4 NA PlanApochromat lens, and an iXon Life EMCCD camera. For monitoring fluorescent marker turn-on, embryos that had been filmed by DIC during early embryogenesis were allowed to develop for 20 additional hours at 20°C before the acquisition of 26 x 1µm z-stack using confocal fluorescence microscopy (488 and 561 nm lasers) and DIC optics. For all measurements, embryos were cropped from time-lapse series and measurements were made using FIJI.^90^ Table S1 has descriptions of the features scored or measured.

### Centriole Positioning in Early Hybrid Embryos

*C. elegans* (OD3701) or *C. brenneri* (LKC28) males were crossed to *C. brenneri* (LKC28) or *C. elegans* (JK574) females and left to mate overnight at 20°C for 24 hours. Female worms were dissected and embryos mounted as described above. Embryos were imaged by collecting 19 x 1.5 µm z-stacks every 30 sec, capturing DIC and fluorescence (561nm laser at 15% power, 2×2 binning, 100ms exposure) through the two-cell stage. FIJI was used to crop and rotate images for scoring. Centrioles were considered detached when centrioles were > 9µm apart. FIJI was used to create maximum projections and to scale images for figures. The image intensities were scaled to best visualize centrosome position within the early embryo.

### Spindle Angle, Pronuclear Localization and Size, and Embryo Aspect Ratio Analysis

Embryos were cropped from timelapse series and measurements were made using FIJI. P0 spindle angle was measured relative to the long axis of the embryo; the angle was assessed from metaphase onset through the following 3.5-4.5 minutes. Images were converted to maximum intensity projections, since centrioles were in different z-planes for part of the first cell division, and then each embryo was scored for centrosome detachment before pronuclear meeting or centrosome mis-localization at pronuclear meeting. Sperm pronucleus length and width was measured at the appearance of the oocyte-derived pronucleus, and sperm-derived pronuclear area was then calculated by using the equation for the area of an ellipse : 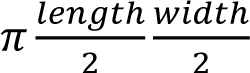. Embryo aspect ratio was calculated as the ratio of cross-sectional width to cross-sectional length of P0 embryos.

### Centriole Positioning and Microtubule tracking

*C. brenneri* (LKC28) females were crossed with *C. brenneri* (LKC28) males or *C. elegans* (OD3701) males, and *C. elegans* (JK574) females were crossed with *C. elegans* (OD3701) males. Female worms were dissected into 250 nM SiR-Tubulin dye (Cytoskeleton, cat# CY-SC002). To generate an agar pad containing SiR-Tubulin, 15µL of 5µM dye was added on top of a 2% agarose pad made in Boyd Buffer. One-cell embryos were transferred by mouth pipette, covered with an 22×22mm coverslip, and sealed with VALAP before imaging. Embryos were imaged on a Nikon Ti2 microscope equipped with a Yokogawa CSU-X1 (Nikon) spinning disk, a 60X, 1.4 NA PlanApochromat lens, and an iXon Life EMCCD camera. A 27 x 1 µm z-stack was collected every min, capturing DIC and fluorescence (561nm at 15% power, 2×2 binning, 100ms exposure, and 640nm at 40% power and 200ms exposure) through the two-cell stage. Images were cropped and rotated for analysis using FIJI. Timelapse SiR-Tubulin sequences were created by generating maximum intensity projections of the best subset of 14 z-slices containing the spindle and were scaled to show the best signal unobscured by the SiR-Tubulin coating the eggshell. Image intensities were scaled independently for each embryo because dye uptake varies between embryos. Centriole channel images were created using maximum projections to best capture the centrioles.

### Developmental Imaging

For the arrest point experiment in Fig. S1A, *C. elegans* males (OD 1719)^50^ or *C. brenneri* males (LKC28) were mated with *C. elegans* females (JK574) or *C. brenneri* females (LKC28) for 24hrs at 20°C. OD1719 animals express germ-layer markers: ectoderm (Pdlg-1::mCherry::his-72 and Pcnd-1::mCherry::his-72), endoderm (Ppha-4::pha-4::GFP), and mesoderm (Phlh-1::his-72::mCherry and Phlh-1::his-72::GFP). For this experiment, embryogenesis was captured by dissecting females in Boyd Buffer and transferring the embryos to a 384-well imaging plate containing 70uL Boyd Buffer. Embryos were imaged over a 10-hour time-course as previously described.^50^

### Immunofluorescence of Early Embryos

Slides for immunofluorescence were generated by dipping in subbing solution prepared by dissolving 0.1g gelatin in 25 mL of distilled water heated to 60°C, cooling to 40°C, adding 0.01g chromalum, and 15mg poly-lysine HBr. Subbing solution was left to stir at 40°C for 2 hours before sterile filtering and storage at 4°C. Slides were dipped in subbing solution heated to 50°C and allowed to dry for 6 hours. 20-30 mated female animals were placed in a 4µl drop of distilled water placed in the center of the slide and an 18×18 coverslip was placed on top. Worms were compressed by pushing on the coverslip with a pipet tip, embryos were pushed out of the mothers, and slides were plunged into liquid nitrogen. Slides were retrieved and a razor blade was used to pop the coverslips off each slide. Slides were immediately immersed in -20°C cold methanol for a 15-minute fixation. Samples were fixed and stained as previously described.^91^ For detection of PAR-2, slides were incubated in unconjugated primary antibody (Mouse-anti-PAR-2 1:1000 dilution) overnight at 4°C, washed, and then incubated with fluorescent secondary antibody (Donkey-anti-Mouse-Cy5) for 30min at room temperature. To stain for centrosomes and microtubules, embryos were incubated with directly-labeled α-tubulin antibodies (DM1-α-FITC 1:1000 dilution; Sigma Aldrich F2168) and anti-γ-tubulin-CY3 (C-terminal antigen: LDEYKAVVQKDYLTRGL; 1:300 dilution)^59^ for 60 minutes at room temperature. Slides were washed with PBST buffer, and 1 μg/ml Hoechst was added during the last 10-minute wash. Two final washes were performed and 15 µl of ProLong Glass Antifade Mount (ThermoFisher) mounting and curing solution was added before covering the embryos with an 18 x 18mm coverslip. The slides were left to cure at room temperature for 24 hours in a dark chamber. Samples were imaged on a DeltaVision (GE Healthcare) epifluorescence scope equipped with a 100X 1.4NA oil immersion objective. Images were deconvolved using SoftWoRx software (Cytvia). Maximum projections were made using FIJI and image intensities adjusted for best visualization of signal.

### Evolutionary analysis of hybrid embryos

*C. brenneri* females were mated to males of *C. sp. 48* and *C. remanei* in a ratio of 1:2 and to males of *C elegans* in a ratio of 1:3 on individual mating plates. *C. sinica* females were mated to *C. remanei* males as described above for *C. brenneri females* and *C. remanei* males. Plates were left overnight at 20°C for mating as described above. Twenty-four hours later, embryos were dissected from gravid female animals in Boyd buffer and mounted as described above. DIC images of early embryogenesis were acquired as described above collecting images every 30s until the 4-cell stage or later. Embryos were cropped and rotated for further analysis. Image names were anonymized by JB, and embryos scored by RG as either a 1 (display phenotype) or 0 (do not display phenotype) for the phenotypes listed in Table S1.

### Divergence time estimates

We downloaded complete *Caenorhabditis* genomes from download.caenorhabditis.org,^89^ extracted the longest isoforms for each protein, and used the species tree estimated by Orthofinder. Branch lengths of the estimated species tree represent molecular phylogenetic distance along the branch.^92^ We used these branch lengths to determine the relative divergence of protein coding sequences between *C. brenneri* and the three other species.

### Sequence Analysis

Protein sequences for analysis were chosen based on membership to the following phenotype categories on Phenobank when targeted by RNA interference: Pronuclear/Nuclear appearance, Centrosome attachment, Asymmetry of Division. In addition, proteins known to be highly conserved such as BEN-1, and proteins of high divergence, such as sperm proteins (SPE-9, SPE-11) were included as points of comparison for divergence scores. The *C. elegans* ortholog for each gene of interest was then used to collect all possible orthologs for *C. brenneri, C. remanei, C. sp. 48*, and *C. sinica* using Orthofinder (Emms 2019) using the default settings. Each ortholog from all species was then locally aligned to the *C. elegans* ortholog from Phenobank using a BLOSUM62 scoring matrix. Orthologs with the highest scores were then used for global pairwise sequence alignment. Alignment scores for each protein were collected in a 2D distance matrix. Distance matrices were then converted to 1D and used to generate heat maps. Only proteins with an identified ortholog for every species analyzed were included in the larger heat map of *C. brenneri* divergence scores. Hierarchical clustering of *C. brenneri* specific divergence scores was generated using R (Rstudio)^93^ (hclust(), method = “complete”) and protein name labels were generated using SimpleMine (https://wormbase.org//tools/mine/simplemine.cgi). All pairwise protein alignments were performed using the Pairwise Aligner tool from Biopython^84^ with a gap score of -12, an open gap score of -0.5 and a maximum of 10,000 possible alignments.

### Meiotic Failure Scoring

Immunofluorescence images of early embryos were scored for meiotic failure by counting polar bodies visible outside of the embryo. Meiotic failures were categorized as follows, meiosis I failure – no polar body visible, meiosis II failure – one polar body visible. DNA capture was scored as positive if chromosomes were in contact with microtubule asters outside of the central spindle.

### P-cell division phenotype

P-cell division timing was scored as before AB cell division, same time as AB cell division, and after AB cell division.

### Quantification and Statistical Analysis

All graphs shown in the manuscript were created and analyzed in R (Rstudio).^93^

---

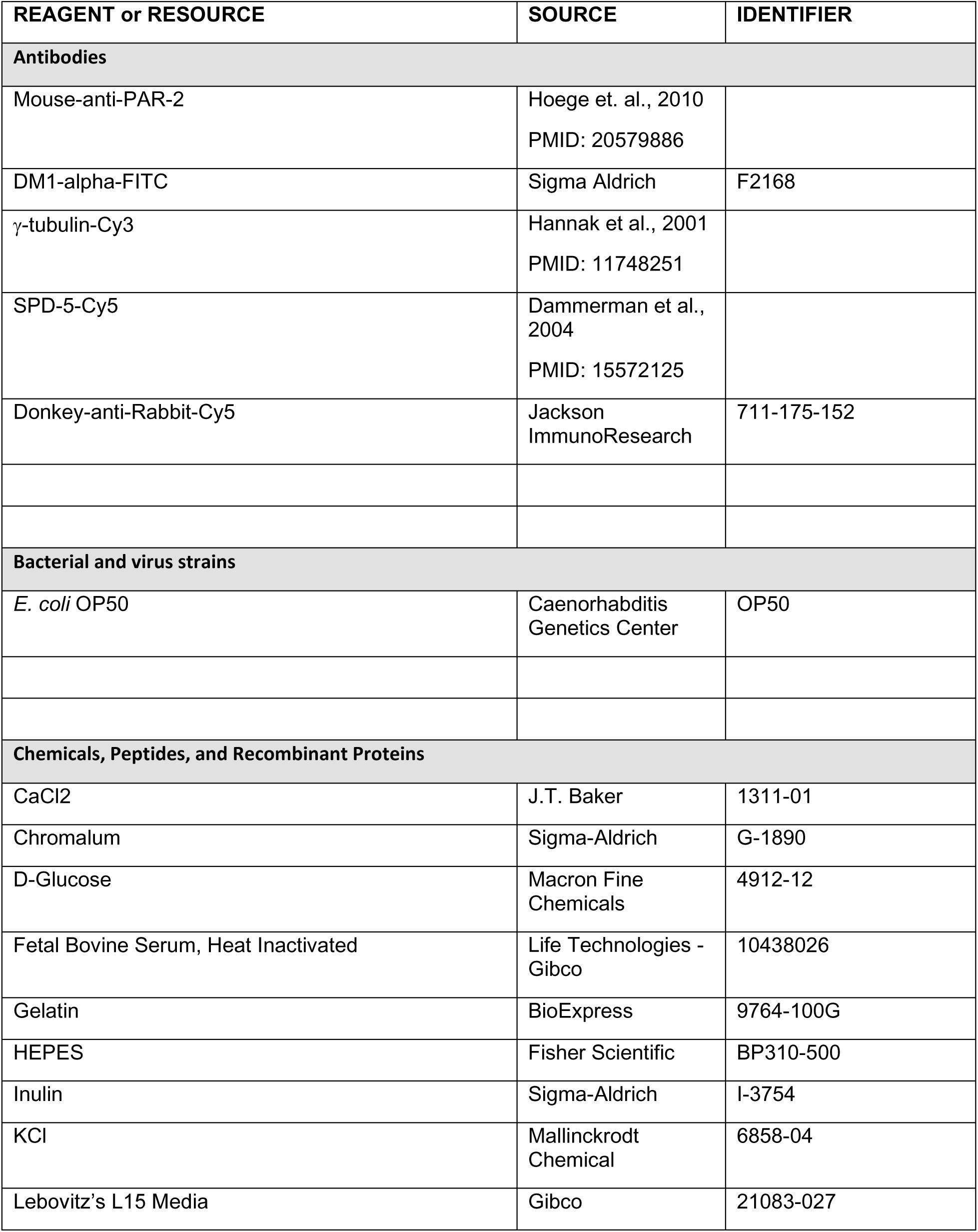
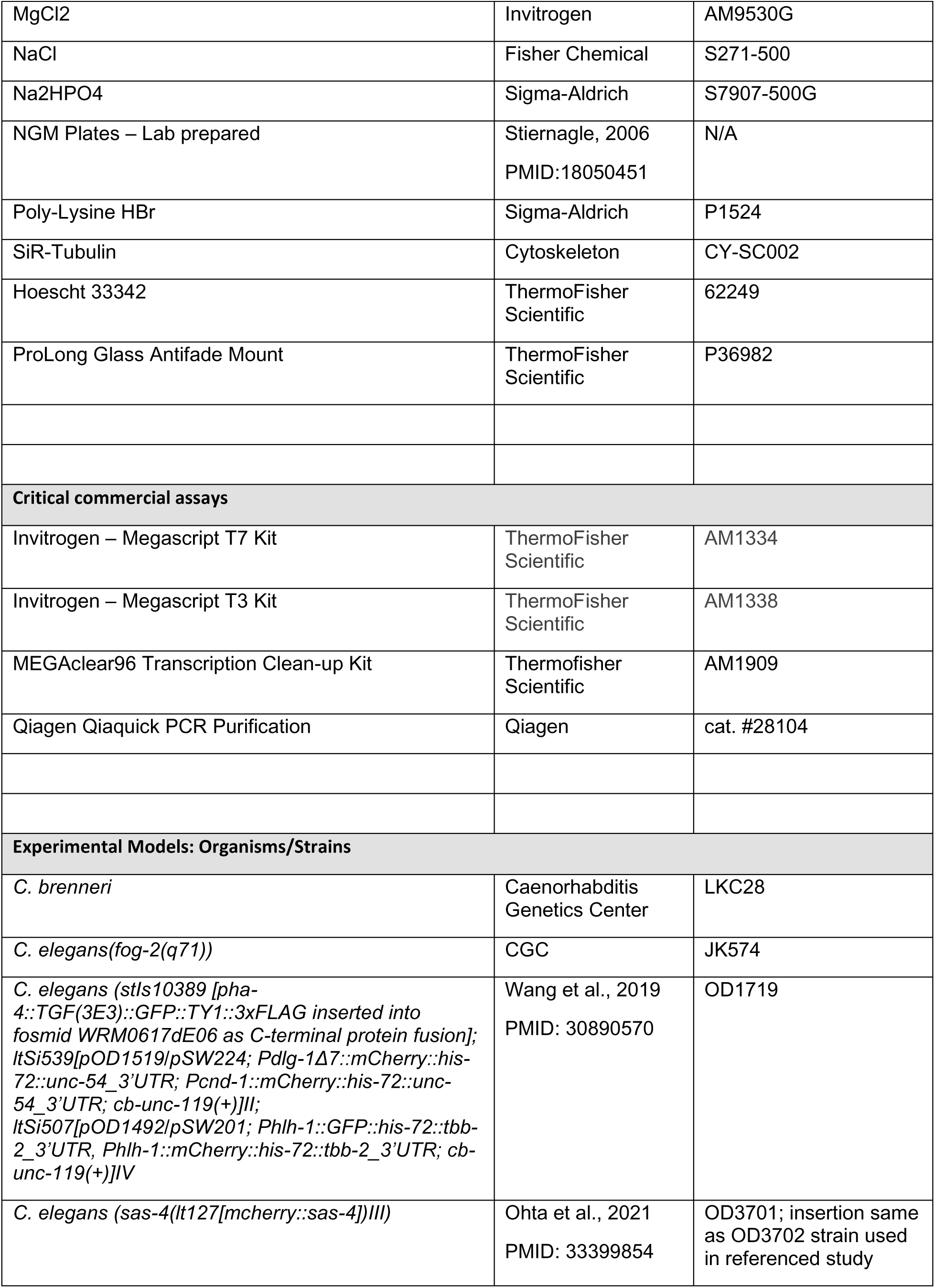
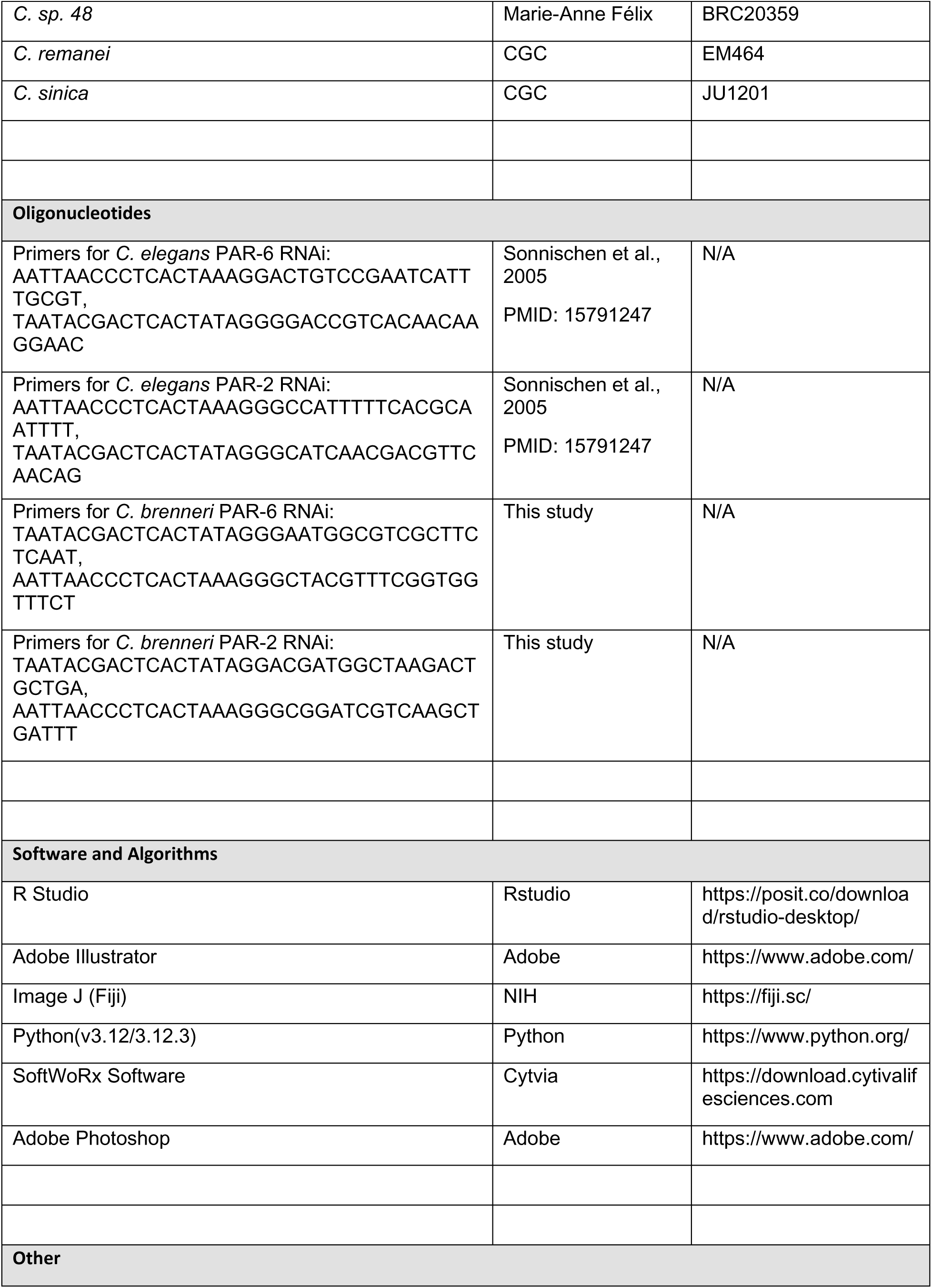
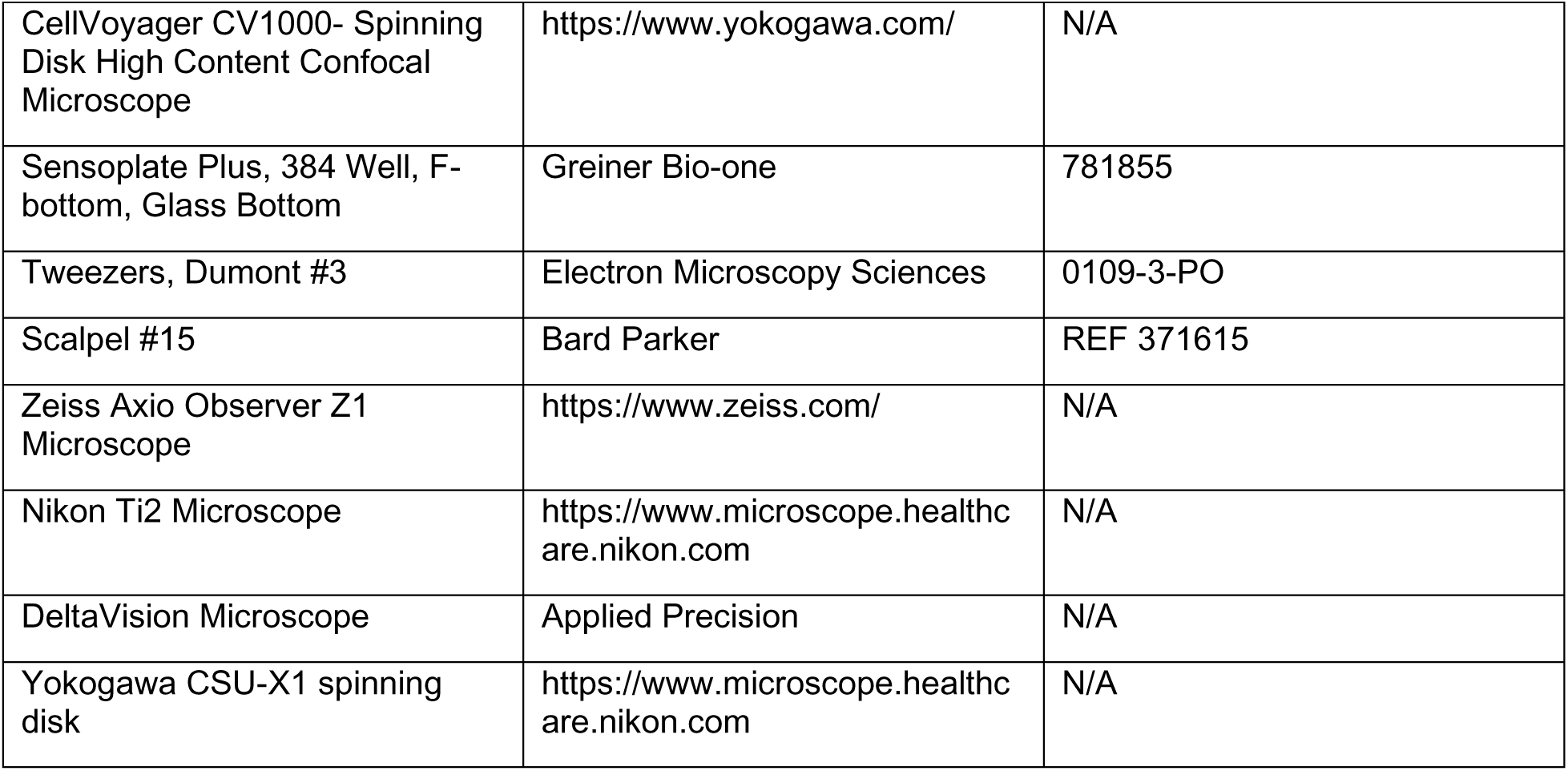
KEY RESOURCES TABLE

