## Supplementary material for "Hybrid incompatibility emerges at the one-cell stage in interspecies *Caenorhabditis* embryos": Key resources table

| REAGENT or RESOURCE | SOURCE | IDENTIFIER |
|---|---|---|
| **Antibodies** | | |
| Mouse-anti-PAR-2 | Hoege et. al., 2010<br>PMID: 20579886 | |
| DM1-alpha-FITC | Sigma Aldrich | F2168 |
| $\gamma$-tubulin-Cy3 | Hannak et al., 2001<br>PMID: 11748251 | |
| SPD-5-Cy5 | Dammerman et al., 2004<br>PMID: 15572125 | |
| Donkey-anti-Rabbit-Cy5 | Jackson ImmunoResearch | 711-175-152 |
| **Bacterial and virus strains** | | |
| *E. coli* OP50 | Caenorhabditis Genetics Center | OP50 |
| **Chemicals, Peptides, and Recombinant Proteins** | | |
| CaCl2 | J.T. Baker | 1311-01 |
| Chromalum | Sigma-Aldrich | G-1890 |
| D-Glucose | Macron Fine Chemicals | 4912-12 |
| Fetal Bovine Serum, Heat Inactivated | Life Technologies - Gibco | 10438026 |
| Gelatin | BioExpress | 9764-100G |
| HEPES | Fisher Scientific | BP310-500 |
| Inulin | Sigma-Aldrich | I-3754 |

| | | |
|---|---|---|
| KCl | Mallinckrodt Chemical | 6858-04 |
| Lebovitz's L15 Media | Gibco | 21083-027 |
| MgCl2 | Invitrogen | AM9530G |
| NaCl | Fisher Chemical | S271-500 |
| Na2HPO4 | Sigma-Aldrich | S7907-500G |
| NGM Plates – Lab prepared | Stiernagle, 2006<br>PMID:18050451 | N/A |
| Poly-Lysine HBr | Sigma-Aldrich | P1524 |
| SiR-Tubulin | Cytoskeleton | CY-SC002 |
| Hoescht 33342 | ThermoFisher Scientific | 62249 |
| ProLong Glass Antifade Mount | ThermoFisher Scientific | P36982 |
| **Critical commercial assays** | | |
| Invitrogen – Megascript T7 Kit | ThermoFisher Scientific | AM1334 |
| Invitrogen – Megascript T3 Kit | ThermoFisher Scientific | AM1338 |
| MEGAclear96 Transcription Clean-up Kit | Thermofisher Scientific | AM1909 |
| Qiagen Qiaquick PCR Purification | Qiagen | cat. #28104 |
| **Experimental Models: Organisms/Strains** | | |
| *C. brenneri* | Caenorhabditis Genetics Center | LKC28 |
| *C. elegans(fog-2(q71))* | CGC | JK574 |
| *C. elegans (stIs10389 [pha-4::TGF(3E3)::GFP::TY1::3xFLAG inserted into fosmid WRM0617dE06 as C-terminal protein fusion]; ltSi539[pOD1519/pSW224; Pdlg-1Δ7::mCherry::his-72::unc-54_3'UTR; Pcnd-* | Wang et al., 2019<br>PMID: 30890570 | OD1719 |

| | | |
|---|---|---|
| *1::mCherry::his-72::unc-54_3'UTR; cb-unc-119(+)]II; ltSi507[pOD1492/pSW201; Phlh-1::GFP::his-72::tbb-2_3'UTR, Phlh-1::mCherry::his-72::tbb-2_3'UTR; cb-unc-119(+)]IV* | | |
| *C. elegans (sas-4(lt127[mcherry::sas-4])III)* | Ohta et al., 2021<br>PMID: 33399854 | OD3701; insertion same as OD3702 strain used in referenced study |
| *C. sp. 48* | Marie-Anne Félix | BRC20359 |
| *C. remanei* | CGC | EM464 |
| *C. sinica* | CGC | JU1201 |
| **Oligonucleotides** | | |
| Primers for *C. elegans* PAR-6 RNAi: AATTAACCCTCACTAAAGGACTGTCCGAATCATTTGCGT, TAATACGACTCACTATAGGGGACCGTCACAACAAGGAAC | Sonnischen et al., 2005<br>PMID: 15791247 | N/A |
| Primers for *C. elegans* PAR-2 RNAi: AATTAACCCTCACTAAAGGGCCATTTTTCACGCAATTTT, TAATACGACTCACTATAGGGCATCAACGACGTTCAACAG | Sonnischen et al., 2005<br>PMID: 15791247 | N/A |
| Primers for *C. brenneri* PAR-6 RNAi: TAATACGACTCACTATAGGGAATGGCGTCGCTTCTCAAT, AATTAACCCTCACTAAAGGGCTACGTTTCGGTGGTTTCT | This study | N/A |
| Primers for *C. brenneri* PAR-2 RNAi: TAATACGACTCACTATAGGACGATGGCTAAGACTGCTGA, AATTAACCCTCACTAAAGGGCGGATCGTCAAGCTGATTT | This study | N/A |
| **Software and Algorithms** | | |
| R Studio | Rstudio | https://posit.co/download/rstudio-desktop/ |

| | | |
|---|---|---|
| Adobe Illustrator | Adobe | https://www.adobe.com/ |
| Image J (Fiji) | NIH | https://fiji.sc/ |
| Python(v3.12/3.12.3) | Python | https://www.python.org/ |
| SoftWoRx Software | Cytvia | https://download.cytivalifesciences.com |
| Adobe Photoshop | Adobe | https://www.adobe.com/ |
| **Other** | | |
| CellVoyager CV1000-Spinning Disk High Content Confocal Microscope | https://www.yokogawa.com/ | N/A |
| Sensoplate Plus, 384 Well, F-bottom, Glass Bottom | Greiner Bio-one | 781855 |
| Tweezers, Dumont #3 | Electron Microscopy Sciences | 0109-3-PO |
| Scalpel #15 | Bard Parker | REF 371615 |
| Zeiss Axio Observer Z1 Microscope | https://www.zeiss.com/ | N/A |
| Nikon Ti2 Microscope | https://www.microscope.healthcare.nikon.com | N/A |
| DeltaVision Microscope | Applied Precision | N/A |
| Yokogawa CSU-X1 spinning disk | https://www.microscope.healthcare.nikon.com | N/A |
